# Urbilaterian origin and evolution of sNPF-type neuropeptide signalling

**DOI:** 10.1101/712687

**Authors:** Luis Alfonso Yañez-Guerra, Xingxing Zhong, Ismail Moghul, Thomas Butts, Cleidiane G. Zampronio, Alexandra M. Jones, Olivier Mirabeau, Maurice R. Elphick

**Affiliations:** Queen Mary University of London, School of Biological & Chemical Sciences, Mile End Road, London, E1 4NS, UK; School of Life Sciences and Proteomics Research Technology Platform, University of Warwick, Coventry, CV4 7AL, UK; Cancer Genetics Unit, Institut Curie, Paris, France

## Abstract

Physiology and behaviour are controlled by neuropeptide signalling systems comprising peptide ligands and cognate receptors. Molecular phylogenetics combined with experimental identification of neuropeptide-receptor pairs has revealed that many neuropeptide signalling systems originated in the urbilaterian common ancestor of protostomes and deuterostomes. Neuropeptide-Y/neuropeptide-F (NPY/NPF)-type signalling is one such example, whereas NPY/NPF-related short-NPF (sNPF)-type signalling has hitherto only been identified in protostomes. Here we report the discovery of a neuropeptide (pQDRSKAMQAERTGQLRRLNPRF-NH_2_) that is the ligand for an sNPF-type receptor in a deuterostome, the starfish *Asterias rubens* (Phylum Echinodermata). Informed by phylogenetic analysis of sequence data, we conclude that the paralogous NPY/NPF-type and sNPF-type signalling systems originated in Urbilateria but NPY/NPF-type signalling was lost in echinoderms. Furthermore, we present evidence that sNPF-type peptides are orthologs of vertebrate prolactin-releasing peptides. Our findings demonstrate the importance of experimental studies on echinoderms for reconstructing the evolutionary history of neuropeptide signalling systems.

## Introduction

Neuropeptides are neuronally secreted signalling molecules that regulate many physiological processes and behaviours in animals, including feeding, digestion, reproduction and social behaviour. They typically exert effects by binding to cognate G-protein coupled receptors (GPCRs) on target cells, which leads to changes in the activity of downstream effectors (e.g. ion channels, enzymes) (Jékely et al. 2018). Investigation of the evolution of neuropeptide signalling has revealed that many of the neuropeptide systems found in vertebrates have orthologs in invertebrate deuterostomes (urochordates, cephalochordates, hemichordates, echinoderms) and protostomes (e.g. arthropods, nematodes, molluscs, annelids, platyhelminthes). Thus, the evolutionary origin of over thirty neuropeptide signalling systems has been traced back to the common ancestor of the Bilateria (Urbilateria) (Mirabeau and Joly 2013; Elphick et al. 2018; Jékely et al. 2018).

One of the neuropeptide signalling systems that originated in Urbilateria is neuropeptide Y (NPY)-type signalling. NPY is a 36-residue peptide that was first isolated from the porcine hypothalamus (Tatemoto et al. 1982; Tatemoto 1982) but that is also expressed by neurons in many other regions of the nervous system (Adrian et al. 1983; Morris 1989) and in peripheral organs such as the gut and cardiovascular system (Holzer et al. 2012; Farzi et al. 2015). Accordingly, NPY is pleiotropic (Pedrazzini et al. 2003), but it is perhaps most widely known as a potent stimulant of food intake in mammals (Minor et al. 2009; Zhang et al. 2011).

NPY belongs to a family of related signalling molecules in vertebrates, including peptide YY (PYY) and pancreatic polypeptide (PP), that evolved from a common ancestral peptide by gene/genome duplication (Larhammar et al. 1993; Larhammar 1996; Elphick et al. 2018). Furthermore, the sequences of NPY-type peptides are highly conserved across the vertebrates, sharing up to 92% identity between mammals and cartilaginous fish (Larhammar et al. 1993; Larhammar 1996; Cerdá-Reverter et al. 2000). A neuropeptide in vertebrates that is evolutionarily related to NPY/PYY/PP-type peptides is prolactin-releasing peptide (PrRP), which was first discovered as a ligand for the orphan receptor hGR3 (Hinuma et al. 1998). Phylogenetic analysis has revealed that PrRP-type receptors are paralogs of NPY/PYY/PP-type receptors and it has been proposed that PrRP-type signalling originated in the vertebrate lineage (Lagerström et al. 2005). However, more recently, orthologs of vertebrate PrRP-type receptors have been identified in invertebrate deuterostomes - the cephalochordate *Branchiostoma floridae* and the hemichordate *Saccoglossus kowalevskii* - indicating that PrRP-type signalling may have originated in a common ancestor of the deuterostomes (Mirabeau and Joly 2013).

An important insight into the evolutionary history of NPY-type peptides was obtained by purification from extracts of a protostome invertebtate, the platyhelminth *Moniezia expansa,* of a peptide immunoreactive with antibodies to the C-terminal hexapeptide of PP (Maule et al. 1991). Sequencing revealed a 39-residue peptide with a similar structure to NPY, but with the C-terminal tyrosine (Y) substituted with a phenylalanine (F). Hence, this invertebrate NPY homolog was named neuropeptide F (NPF) (Maule et al. 1991). Subsequently, NPF-type neuropeptides have been identified in other protostomian invertebrates, including other platyhelminths (Curry et al. 1992), molluscs (Leung et al. 1992; Rajpara et al. 1992), annelids (Díaz-Miranda et al. 1991; Veenstra 2011; Conzelmann et al. 2013; Bauknecht and Jékely 2015) and arthropods (Brown et al. 1999), and these peptides typically have a conserved C-terminal RPRFamide motif and range in length from 36 to 43 residues.

Following the discovery of *M. expansa* NPF, antibodies to this peptide were generated and used to assay for related peptides in other invertebrates. Interestingly, this resulted in the discovery of two novel peptides, ARGPQLRLRFamide and APSLRLRFamide, in brain extracts from the Colorado potato beetle *Leptinotarsa decemlineata* (Spittaels et al. 1996). As these peptides were isolated using antibodies to *M. expansa* NPF, they were originally referred to as NPF-related peptides. However, because they are much shorter in length than NPF, they were later renamed as short neuropeptide F (sNPF) (Vanden Broeck 2001) and homologs were identified in other insects (Schoofs et al. 2001). Furthermore, alignment of NPY-type peptides and precursors from vertebrates with NPF-type and sNPF-type peptides and precursors from protostomes revealed that whilst NPF peptides are clearly closely related (orthologous) to vertebrate NPY peptides, sNPF peptides and precursors exhibit too many differences to be considered orthologs of NPY/NPF-type peptides and precursors (Nässel and Wegener 2011). Further evidence that chordate NPY-type and invertebrate NPF-type neuropeptides are orthologous has been provided by similarity-based clustering methods, showing that the NPY-type and NPF-type precursors form a pan-bilaterian cluster, whereas sNPF-type precursors form a separate cluster (Jékely 2013). Thus, sNPF-type peptides are considered to be a family of neuropeptides that is distinct from the NPY/NPF-type family of neuropeptides.

A receptor for sNPF-type peptides was first identified in the fruit fly *Drosophila melanogaster* with the deorphanisation of the G-protein coupled receptor CG7395 (Mertens et al. 2002), which was previously annotated as a homolog of mammalian NPY-type receptors. Subsequently, sNPF receptors have been identified in other insects, including the fire ant *Solenopsis Invicta* (Chen and Pietrantonio 2006), the mosquitoes *Anopheles gambiae* (Garczynski et al. 2007) and *Aedes aegypti* (Christ et al. 2018), the desert locust *Schistocerca gregaria* (Dillen, Zels, et al. 2013), the oriental fruit fly *Bactrocera dorsalis* (Jiang et al. 2017) and in the silkworm *Bombyx mori* (Yamanaka et al. 2008; Ma et al. 2017).

A variety of physiological roles have been attributed to sNPF-type peptides in insects, with the most consistent being actions related to the regulation of feeding behaviour. For example, in *D. melanogaster* overexpression of sNPF increases food intake both in larvae and adults, whilst loss-of-function sNPF-mutants show reduced food intake (Lee et al. 2004). In *A. mellifera*, food-deprivation upregulates transcription of the sNPF receptor gene and quantitative peptidomics reveals a correlation between sNPF peptide levels and the pollen foraging predisposition of worker bees (Brockmann et al. 2009; Ament et al. 2011). In contrast, in the locust *S. gregaria* sNPF inhibits food intake, whilst knockdown of sNPF precursor or sNPF receptor gene expression significantly increases food intake, indicating that sNPF acts as a satiety signal in locusts (Dillen, Verdonck, et al. 2013; Dillen, Zels, et al. 2013). In the cockroach *Periplaneta americana*, starvation followed by feeding increases and then decreases, respectively, the number of sNPF-immunoreactive cells in the midgut epithelium.

Since sNPF signalling was discovered in insects, it was initially thought that this neuropeptide system may be unique to arthropods (Nässel and Wegener 2011). However, a large-scale phylogenetic analysis of G-protein coupled neuropeptide receptors revealed that sNPF-type signalling may also be present in other protostomes (Mirabeau and Joly 2013). Thus, an expanded family of neuropeptide receptors in the nematode *C. elegans* that had originally been annotated as NPY/NPF-type receptors (Cardoso et al. 2012) were found to be orthologs of insect sNPF-receptors (Mirabeau and Joly 2013). Furthermore, whilst NPY/NPF-type peptides and their receptors were identified as a bilaterian neuropeptide signalling system, it was proposed that sNPF-type signalling may be restricted to protostomes (Mirabeau and Joly 2013). Subsequently, sNPF-type peptides and a cognate receptor have been characterised in the bivalve mollusc *Crassostrea gigas*, confirming the occurrence of this signalling system in the lophotrochozoan branch of the protostomes (Bigot et al. 2014). Furthermore, functional studies revealed that starvation causes upregulated expression of the sNPF precursor and receptor in the visceral ganglia of *C. gigas*, providing evidence of an evolutionarily conserved role of sNPF-type peptides in feeding-related processes in protostomes (Bigot et al. 2014). Orthologs of the *C. gigas* sNPF-type peptides have been identified in other molluscs, and interestingly other functions of this family of neuropeptides have been reported. For example, in the gastropod *Lymnaea stagnalis* sNPF-type peptides are upregulated in response to infection with the parasite *Trichobilharzia ocellata*, causing an accelerated increase in body weight, reduced metabolism and retarded development of the reproductive system (Hoek et al. 2005). Furthermore, in the cephalopod *Sepia officinalis* the sNPF-type peptide GNLFRFamide acts as a myoactive peptide on the rectum, increasing the frequency, tonus and amplitude of the rectal contractions (Zatylny-Gaudin et al. 2010).

Important insights into neuropeptide evolution have been obtained recently by pharmacological characterisation of G-protein coupled neuropeptide receptors in invertebrate deuterostomes (Kawada et al. 2010; Jékely 2013; Mirabeau and Joly 2013; Elphick and Mirabeau 2014; Roch et al. 2014; Satoh et al. 2014; Semmens et al. 2016; Tian et al. 2016; Yañez-Guerra et al. 2018). However, currently little is known about the occurrence and characteristics of NPY/NPF/sNPF-related signalling systems in invertebrate deuterostomes. Phylogenetic analysis of bilaterian G-protein coupled neuropeptide receptors has demonstrated the occurrence of NPY/NPF receptor-related proteins in ambulacrarians – the echinoderm *Strongylocentrotus purpuratus* and the hemichordate *Saccoglossus kowalevskii* (Mirabeau and Joly 2013). Furthermore, the precursor of a putative NPY/NPF-type peptide was identified in *S. kowalevskii* (Mirabeau and Joly 2013; Elphick and Mirabeau 2014). A candidate NPY/NPF-type precursor has also been identified in the cephalochordate *Branchiostoma floridae*, but an NPY/NPF-type receptor has yet to be identified in this species (Mirabeau and Joly 2013; Elphick and Mirabeau 2014). A more recent finding was the discovery of a family neuropeptide precursor-type proteins in echinoderms that contain a peptide that shares sequence similarity with NPY/NPF-type peptides (Zandawala et al. 2017). However, it is not known if these proteins are orthologs of vertebrate NPY-type precursor peptides and protostome NPF-type precursor peptides. To address this issue, detailed analysis of the sequences of the echinoderm NPY/NPF-like peptides and precursors and the genes encoding these peptides/proteins is needed. Furthermore, the receptors for echinoderm NPY/NPF-like peptides need to be identified. Accordingly, here we report the biochemical and pharmacological characterisation of a NPY/NPF/sNPF-related signalling system in an echinoderm – the starfish *Asterias rubens*. Furthermore, informed by detailed phylogenetic analyses, we provide new insights into the evolutionary history of NPY/NPF-type and sNPF-type signalling in the Bilateria.

## Results

### NPY-like peptides in the starfish *Asterias rubens* and in other echinoderms

The sequence of a transcript (contig 1060225; accession number MK033631.1) encoding the precursor of an NPY-like neuropeptide has been reported previously based on analysis of *A. rubens* neural transcriptome sequence data (Zandawala et al. 2017). Here, a cDNA encoding this precursor was cloned and sequenced, revealing that the open reading frame encodes a 108-residue protein comprising a predicted 19-residue signal peptide, a 23-residue NPY-like peptide sequence with an N-terminal glutamine residue and a C-terminal glycine residue, followed by a putative monobasic cleavage site (Supplementary Figure 1A). Analysis of radial nerve cord extracts using mass spectrometry (LC-MS-MS) revealed the presence of a peptide with the structure pQDRSKAMQAERTGQLRRLNPRF-NH_2_, showing that the N-terminal glutamine and C-terminal glycine in the precursor peptide are post-translationally converted to a pyroglutamate residue and amide, respectively (Supplementary Figure 1B). Having determined the structure of this peptide, we provisionally named it *A. rubens* NPY-like peptide or ArNPYLP.

The ArNPYLP sequence was aligned with related peptides from other echinoderms and with NPY/NPF-type peptides from other phyla (Figure 1). This revealed that ArNPYLP and a closely related peptide in the starfish *Acanthaster planci* both share a C-terminal PRFamide sequence with several protostome NPF-type peptides. In contrast, related peptides in two other echinoderms, a brittle star and a sea urchin, have a C-terminal RYamide motif, which is a characteristic of vertebrate NPY-type peptides. However, the alignment also revealed that the echinoderm NPY-like peptides are shorter (22-25 residues) than NPY/NPF-type peptides in other taxa (30-41 residues). Furthermore, the echinoderm peptides lack two proline (P) residues that are a conserved feature of the N-terminal region of many NPY/NPF-type peptides, with the exception some peptides that have only one of these proline residues and a peptide in the cephalochordate *Branchiostoma floridae* that has neither (Figure 1). Furthermore, there are four other residues that are highly conserved in bilaterian NPY/NPF peptides - tyrosine (Y), leucine (L), tyrosine (Y) and isoleucine (I) residues, which are marked with asterisks in Figure 1. Importantly, none of these residues are present in the echinoderm NPY-like peptides. It is noteworthy, however, that all but one of the aforementioned six conserved residues in NPY/NPF-type peptides are present in a peptide from a species belonging to a sister phylum of the echinoderms – the hemichordate *Saccoglossus kowalevskii* (Figure 1) (Mirabeau and Joly 2013; Elphick and Mirabeau 2014). Collectively these findings indicated that ArNPYLP and related peptides in other echinoderms may not be orthologs of NPY/NPF-type peptides.

**Figure 1.**
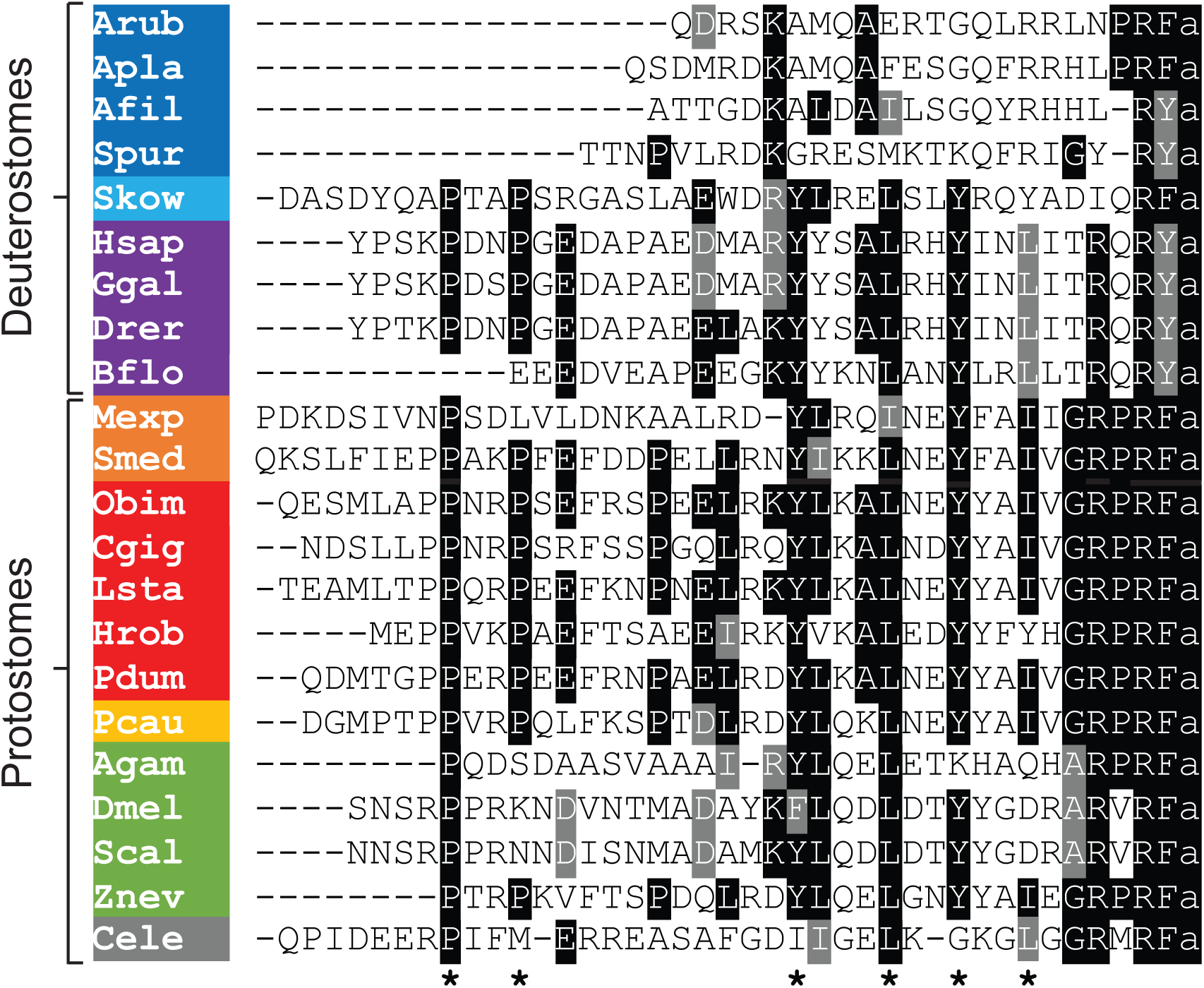
Comparison of the sequences of echinoderm NPY-like peptides with NPY/NPF-type peptides from other taxa. Conserved residues are highlighted in black or grey. The asterisks indicate residues that have been shown to be important for the three-dimensional structure of the NPY/NPF-type peptides, but which are not present in the echinoderm NPY-like peptides. Species names are highlighted in phylum-specific or superphylum-specific colours: dark blue (Echinodermata), light blue (Hemichordata), purple (Chordata), orange (Platyhelminthes), red (Lophotrochozoa), yellow (Priapulida), green (Arthropoda), grey (Nematoda). Species names are as follows: Arub (*Asterias rubens*), Apla (*Acanthaster planci*), Afil (*Amphiura filiformis*), Spur (*Strongylocentrotus purpuratus*), Skow (*Saccoglossus kowalevskii*), Hsap (*Homo sapiens*), Ggal (*Gallus gallus*), Drer (*Danio rerio*), Bflo (*Branchiostoma floridae*), Mexp (*Moniezia expansa*), Smed (*Schmidtea mediterranea*), Obim (*Octopus bimaculoides*), Cgig (*Crassostrea gigas*), Lsta (*Lymnaea stagnalis*), Hrob (*Helobdella robusta*), Pdum (*Platynereis dumerilii*), Pcau (*Priapulus caudatus*), Agam (*Anopheles gambiae*), Dmel (*Drosophila melanogaster*), Scal (*Stomoxys calcitrans*), Znev (*Zootermopsis nevadensis*), Cele (*Caenorhabditis elegans*). The accession numbers of the sequences included in this alignment are listed in supplementary table 1.

### The exon-intron structure of echinoderm NPYLP genes is different to NPY/NPF genes

To investigate further our proposition that echinoderm NPY-like neuropeptides may not be orthologs of NPY/NPF-type neuropeptides, we compared the exon-intron structure of genes encoding these peptides. Previous studies have reported that a conserved feature of NPY/NPF genes is an intron that interrupts the coding sequence for NPY/NPF-type peptides, with the intron located between the second and third nucleotide of the codon for the arginine residue of the C-terminal RF or RY dipeptide (Mair et al. 2000). Here we show this conserved feature in NPY/NPF genes in species from several animal phyla, including a hemichordate (sister phylum to the echinoderms), chordates, molluscs, an annelid, a priapulid, an arthropod and a nematode (Figure 2). Because genome sequence data are currently not available for the starfish *A. rubens*, we examined the structure of the NPYLP gene in echinoderm species where genome sequences have been obtained – the starfish *A. planci* and the sea urchin *S. purpuratus*. This revealed that in the echinoderm NPYLP genes the coding sequence for NPYLP is interrupted by an intron, but it is located in a different position to the intron that interrupts the coding sequence for NPY/NPF-type peptides. Thus, it does not interrupt the codon for the arginine of the C-terminal RF or RY motif, but instead it is located between the first and second nucleotide of the codon for an alanine (*A. planci*) or glycine (*S. purpuratus*) residue located in the N-terminal or central regions, respectively, of the NPYLPs (Figure 2). Another difference is that typically in NPY/NPF genes there is another intron that interrupts the coding sequence in the C-terminal region of the precursor protein, whereas in the echinoderm NPYLP genes the coding sequence for the C-terminal region of the precursor protein is not interrupted by an intron. Collectively, these findings provide further evidence that the echinoderm NPY-like peptides are not orthologs of NPY/NPF-type neuropeptides.

**Figure 2.**
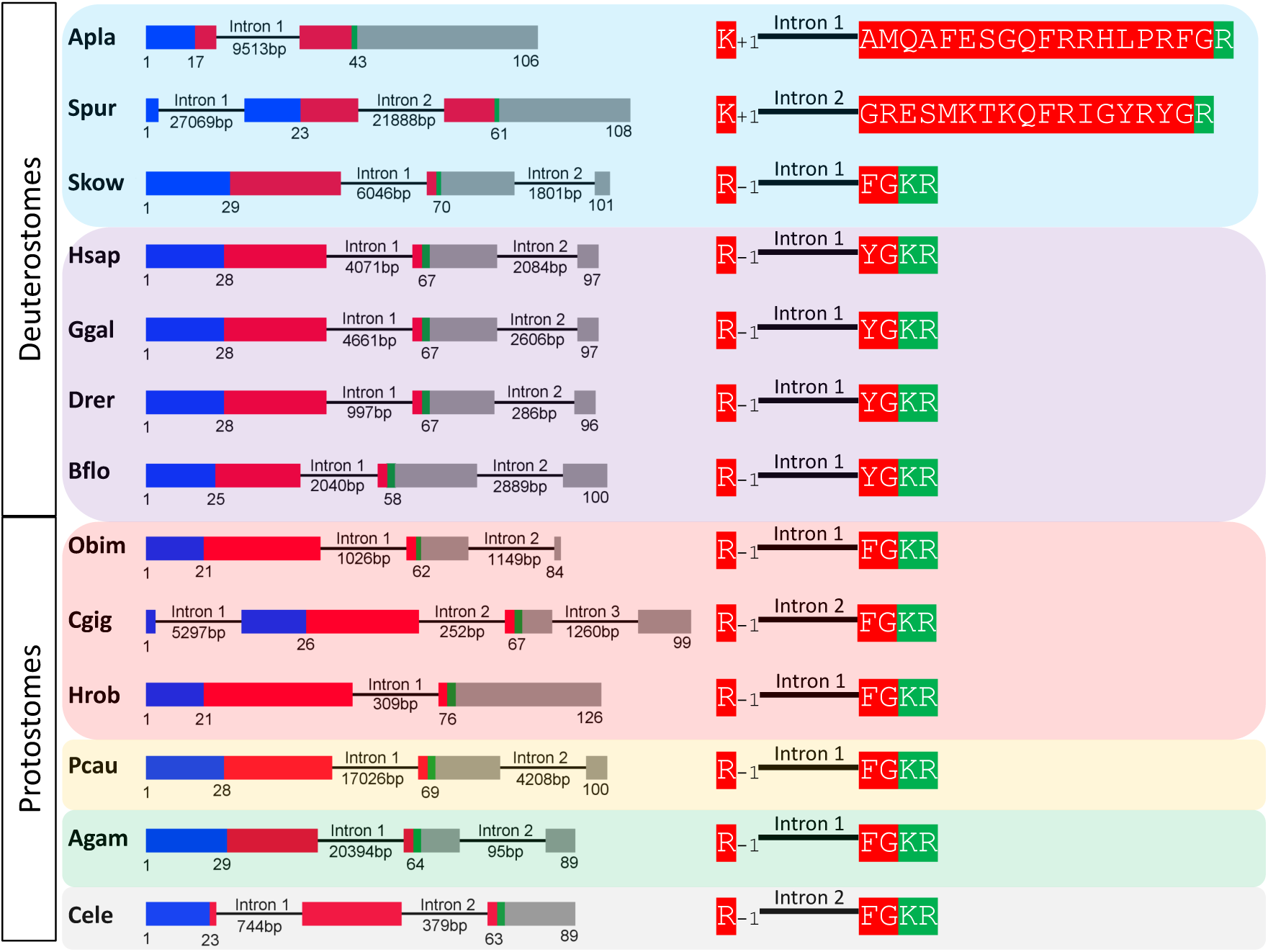
Comparison of the exon/intron structure of genes encoding echinoderm precursors of NPY-like peptides and genes encoding NPY/NPF-type precursors in other taxa. The left side of the figure shows schematic representations of the gene structures, with protein-coding exons shown as rectangles and introns shown as lines (with intron length stated underneath). The protein-coding exons are colour-coded to show regions that encode the N-terminal signal peptide (blue), the neuropeptide (red), monobasic or dibasic cleavage sites (green) and other regions of the precursor protein (grey). The right side of the figure shows how an intron interrupts the coding sequence of the neuropeptides, but the position of the intron is different for echinoderm NPY-like peptides and bilaterian NPY/NPF-type neuropeptides. In genes encoding echinoderm NPY-like peptides the intron interrupts the coding sequence in the N-terminal or central region of the NPY-like peptide, with the intron located between the first and second nucleotides of the codon for an alanine (*A. planci*) or glycine (*S. purpuratus*) residue and the position of the intron in the reading frame is therefore represented as +1. In contrast, in bilaterian NPY/NPF genes the intron interrupts the coding sequence for NPY/NPF-type peptides between the second and third nucleotide of the codon for the arginine residue of the C-terminal RF or RY dipeptide and the position of the intron in the reading frame is therefore represented as −1. Species names are as follows: Apla *(Acanthaster planci),* Spur (*Strongylocentrotus purpuratus),* Skow (*Saccoglossus kowalevskii*), Hsap (*Homo sapiens*), Ggal (*Gallus gallus),* Drer (*Danio rerio),* Bflo (*Branchiostoma floridae),* Obim (*Octopus bimaculoides*), Cgig (*Crassostrea gigas*), Hrob (*Helobdella robusta*), Pcau (*Priapulus caudatus),* Agam (*Anopheles gambiae)*, Cele *(Caenorhabditis elegans).* The accession numbers for the sequences of the precursors shown in this figure are listed in supplementary table 2.

### Discovery of orthologs of sNPF-type receptors in *A. rubens* and other echinoderms

Having obtained evidence that the echinoderm NPY-like peptides are not orthologs of NPY/NPF-type neuropeptides in other bilaterians, we then investigated the occurrence in *A. rubens* and other echinoderms of proteins related to G-protein coupled receptors that mediate effects of NPY/NPF-type peptides and sNPF-type peptides in other bilaterians. Using receptor sequences for the *H. sapiens* NPY-type, *D. melanogaster* NPF-type and *D. melanogaster* sNPF-type receptors as queries for similarity-based analysis of *A. rubens* neural transcriptome sequence data, a transcript (contig 1120879) encoding a 386-residue protein was identified as the best hit (Supplementary Figure 2). Homologs of this protein were also identified in other echinoderms, including the starfish *A. planci*, the sea urchin *S. purpuratus* and the sea cucumber *A. japonicus*. To determine the relationship of these echinoderm receptors with other bilaterian neuropeptide receptors, we performed a phylogenetic analysis using the maximum likelihood method. For this analysis, in addition to bilaterian NPY/NPF-type receptors and protostome sNPF-type receptors, we also included receptors that are closely related to NPY/NPF-type and sNPF-type receptors - prolactin-releasing peptide-type, GPR83-type, tachykinin (TK)-type and luqin (LQ)-type receptors. This revealed that the echinoderm receptors are positioned within a branch of the phylogenetic tree that comprises NPY/NPF-type and sNPF-type receptors, with the other receptor types included in the analysis occupying an outgroup position (Figure 3). Furthermore, the echinoderm receptors are not positioned in a clade comprising NPY/NPF-type receptors but instead they are positioned in a clade comprising sNPF-type receptors, with bootstrap support of >90 %. Thus, we conclude that the echinoderm receptors are orthologs of protostome sNPF-type receptors and accordingly we named the *A. rubens* receptor Ar-sNPFR. Furthermore, we hypothesised that this protein may be the receptor for the *A. rubens* peptide that we have referred to hitherto as ArNPYLP.

**Figure 3.**
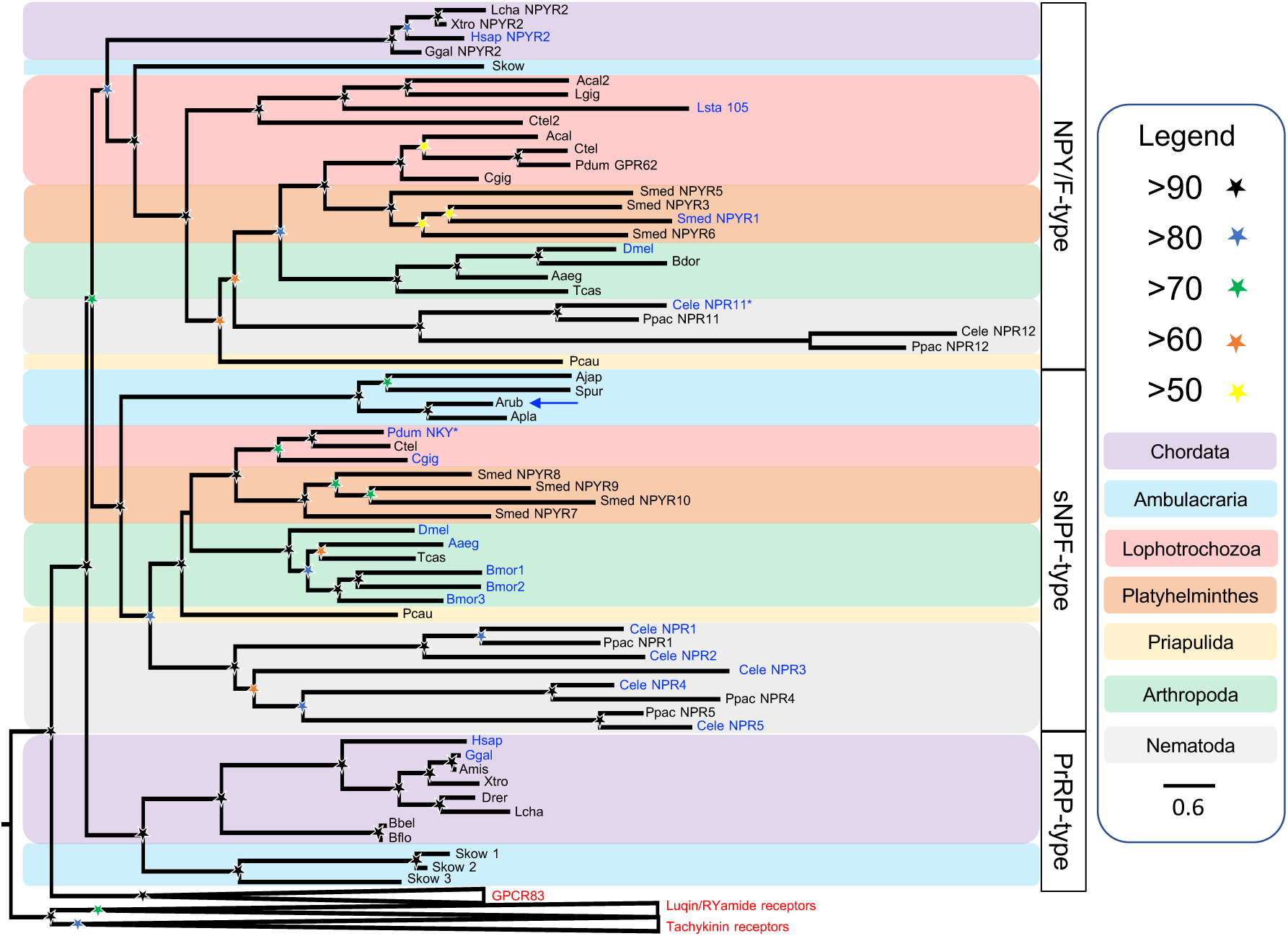
Phylogenetic tree showing that echinoderm NPY-like receptor proteins are not orthologs of bilaterian NPY/NPF-type receptors but are orthologs of protostome sNPF-type receptors. The tree, which was generated in W-IQ-tree 1.0 using the Maximum likelihood method, comprises two distinct clades: a NPY/NPF-type receptor clade and a sNPF-type receptor clade. Also included in the tree are prolactin-releasing peptide-type receptors, which are paralogs of NPY-type receptors in vertebrates. GPR83-type, luqin-type, and tachykinin-type receptors were included as outgroups to root the tree. The stars represent bootstrap support (1000 replicates, see legend) and the coloured backgrounds represent different taxonomic groups, as shown in the key. The names with text in blue represent the receptors in which ligands have been experimentally confirmed. The asterisks highlight receptors where the reported ligand is atypical when compared with ligands for receptors in the same clade. The *Asterias rubens* receptor that was characterised in this study (see Figure 4) is labelled with a blue arrow. Species names are as follows: Aaeg (*Aedes aegypti*), Acal *(Aplysia californica)*, Ajap (*Apostichopus japonicus*), Amis (*Alligator mississippiensis*), Apla (*Acanthaster planci*), Arub *(Asterias rubens)*, Bbel (*Branchiostoma belcheri*), Bdor (*Bactrocera dorsalis*), Bflo (*Branchiostoma floridae*), Bmor (*Bombyx mori*), Cele *(Caenorhabditis elegans)*, Cgig (*Crassostrea gigas*), Ctel *(Capitella teleta)*, Dmel *(Drosophila melanogaster)*, Drer (Danio rerio), Ggal (*Gallus gallus*), Hsap (*Homo sapiens*), Lcha (*Latimeria chalumnae*), Lgig *(Lottia gigantea)*, Lsta (*Lymnaea stagnalis*), Pcau (*Priapulus caudatus*), Pdum (*Platynereis dumerilii*), Ppac (*Pristionchus pacificus*), Skow (*Saccoglossus kowalevskii*), Smed (*Schmidtea mediterranea*), Spur (*Strongylocentrotus purpuratus*), Tcas *(Tribolium castaneum),* Xtro *(Xenopus tropicalis)*. The accession numbers of the sequences used for this phylogenetic tree are listed in supplementary table 3.

### Pharmacological characterisation of Ar-sNPFR

Having identified Ar-sNPFR as a candidate receptor for ArNPYLP, a cDNA encoding this receptor was cloned and sequenced (Supplementary Figure 2) and its sequence has been deposited in GenBank under accession number MH807444.1. Analysis of the sequence of Ar-sNPFR using Protter, revealed seven predicted transmembrane domains, as expected for a G-protein coupled receptor (Supplementary Figure 3). The cloned receptor was then co-expressed with Gα16 in CHO-K1 cells expressing apoaequorin to produce the cell system CHO-Ar-sNPFR. Synthetic ArNPYLP (pQDRSKAMQAERTGQLRRLNPRF-NH_2_) was then tested as a candidate ligand for Ar-sNPFR at concentrations ranging from 10^−14^ M to 10^−5^ M, comparing with cells incubated in assay media without the addition of the peptide. This revealed that ArNPYLP at a concentration of 10^−5^ M triggers luminescence responses (defined as 100%) in CHO-Ar-sNPFR cells that were approximately five times the background luminescence detected with the assay media used to dissolve the peptide (Figure 4A), demonstrating that ArNPYLP acts as a ligand for the receptor. Furthermore, ArNPYLP induced dose-dependent luminescence in CHO-Ar-sNPFR cells with a half-maximal response concentration (EC_50_) of 1.5 × 10^−10^ M (Figure 4B). Importantly, no response to ArNPYLP was observed in CHO-K1 cells transfected with the vector alone, demonstrating that the signal observed in CHO-Ar-sNPFR cells exposed to ArNPYLP can be attributed to activation of the transfected receptor (Supplementary Figure 4). Because ArNPYLP contains a potential dibasic cleavage site (see underlined arginine residues in its sequence: pQDRSKAMQAERTGQL<u>RR</u>LNPRF-NH_2_), we hypothesised that the C-terminal pentapeptide of ArNPYLP (LNPRFamide) may also be generated from ArNPYLP *in vivo*. Therefore, we also tested synthetic LNPRFamide as a candidate ligand for Ar-sNPFR. However, this peptide did not induce luminescence responses in CHO-Ar-sNPFR cells (Figure 4B). So we conclude that the 22-residue amidated peptide ArNPYLP is the natural ligand for Ar-sNPFR in *A. rubens*. Furthermore, on this basis we changed the name of this peptide from ArNPYLP to Ar-sNPF.

**Figure 4.**
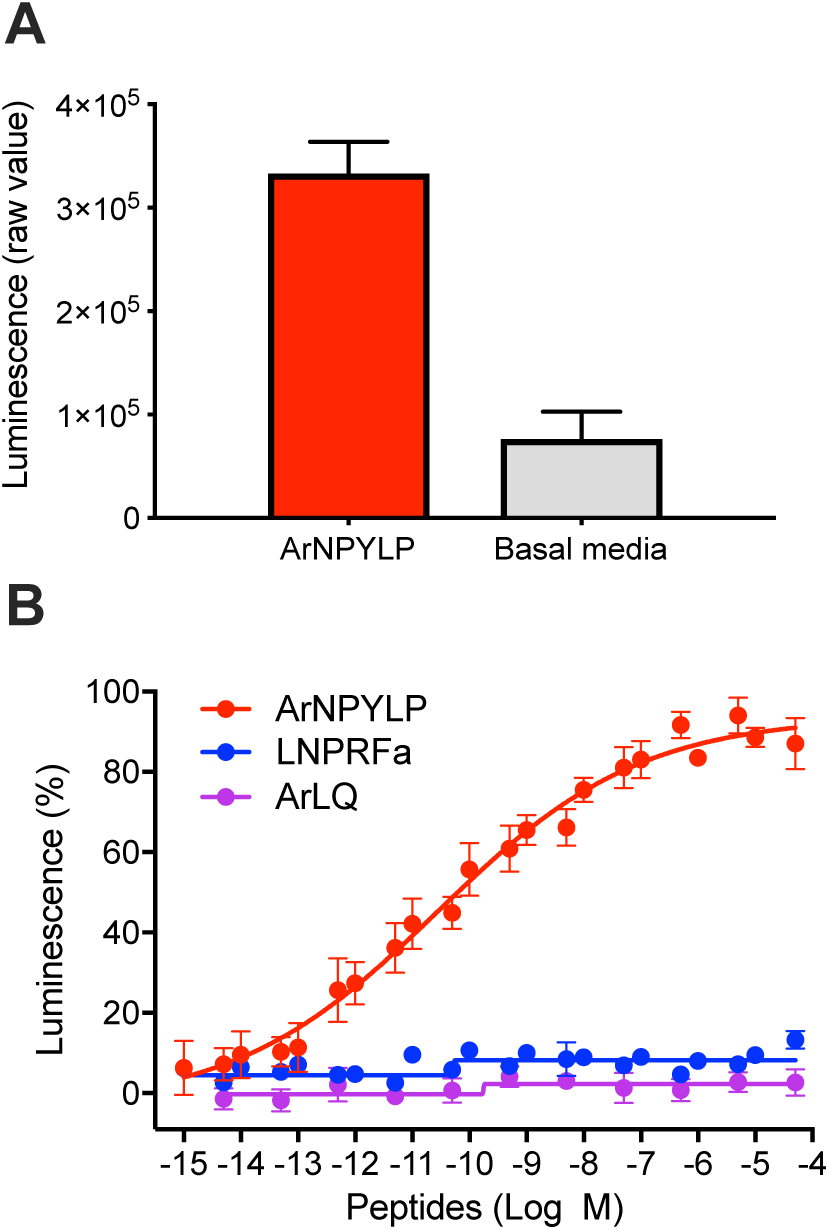
*A. rubens* NPY-like peptide acts as a ligand for a sNPF-type receptor. (**A**) The *A. rubens* NPY-like peptide ArNPYLP (10^-5^ M; red bar) triggers luminescence in CHO-K1 cells expressing the *A. rubens* sNPF-type receptor Ar-sNPFR, the promiscuous G-protein G_α16_ and the calcium-sensitive luminescent GFP-apoaequorin fusion protein G5A. For comparison, the background luminescence of cells that were not exposed to ArNPYLP is shown (basal media; grey bar). Mean values (± S.E.M) were determined from three independent experiments performed in triplicate (**B**). Graph showing the selectivity of Ar-sNPFR as a receptor for ArNPYLP. ArNPYLP causes dose-dependent luminescence in CHO-K1 cells expressing Ar-sNPFR, with an EC_50_ of 0.15 nM. The receptor is not activated by a C-terminal pentapeptide fragment of ArNPYLP (LNPRFamide) or by the *A. rubens* luqin-type peptide ArLQ. Each point represents mean values (± S.E.M) from at least three independent experiments done in triplicate.

### Comparison of the sequences of Ar-sNPF and orthologs in other echinoderms with sNPF-type peptides from other taxa

Having identified the 22-residue amidated peptide pQDRSKAMQAERTGQL<u>RR</u>LNPRF-NH_2_ (Ar-sNPF) as the ligand for Ar-sNPFR, it was of interest to compare the sequences of Ar-sNPF and echinoderm orthologs of this peptide with sNPF-type peptides that have been identified in protostome phyla. Therefore, we aligned the echinoderm peptides with: i). sNPF-type peptides that have been identified in insects (Mertens et al. 2002; Yamanaka et al. 2008; Ma et al. 2017) ii). peptides derived from the *Caenorhabidits elegans* FLP-15, FLP-18 and FLP-21 precursor proteins, which are ligands for sNPF-type receptors in this species (Kubiak, Larsen, Zantello, et al. 2003; Rogers et al. 2003; Kubiak et al. 2008; Cohen et al. 2009; Ezcurra et al. 2016), iii). peptides derived from the *C. elegans* FLP-3 precursor proteins, which share sequence similarity with peptides derived from the *C. elegans* FLP-15, FLP-18 and FLP-21 precursor proteins and iv). peptides that have been shown to be ligands for a sNPF-type receptor in the mollusc *Crassostrea gigas* (Bigot et al. 2014) and orthologous peptides in other molluscan, annelid and platyhelminth species (Figure 5). This revealed sequence similarities shared between the echinoderm peptides and sNPF-type peptides in other phyla. For example, the echinoderm peptides typically have a serine-glycine (SG) motif (or TG in Ar-sNPF, which represents a conservative substitution) in their central region and this aligns with an N-terminal SG motif in a sNPF-type peptide from the insect *Tribolium castaneum* and with a serine or glycine residue in the N-terminal region of other protostome sNPF-type peptides (Figure 5). Furthermore, the C-terminal region of the echinoderm peptides also shares sequence similarity with the C-terminal region of protostome sNPF-type peptides. Thus, Ar-sNPF has the C-terminal sequence LNPRFamide and likewise sNPF-type peptides with a C-terminal LxxRFamide motif occur in some insect species and sNPF-type peptides with a C-terminal LxRFamide or LxRYamide (with Y being a conservative substitution) occur in some molluscan and annelid species. There are, however, also notable differences between the echinoderm peptides and the protostome sNPF-type peptides. Thus, in addition to obvious differences in peptide length, many protostome sNPF-type peptides have a conserved proline residue but this is not a feature of the echinoderm peptides (Figure 5). Finally, a noteworthy highly variable feature of protostome sNPF-type precursors is the number of neuropeptides they give rise to. Thus, the echinoderm precursors contain a single neuropeptide, whereas the number of sNPF-type peptides derived from protostome precursors range from one (*C. elegans* FLP-21, *T. castaneum*), to three or four (e.g. *S. mediterranea*, *D. melanogaster*, *C. gigas*) to as many as seven (e.g. *C. elegans* FLP3, *P. dumerilii*).

**Figure 5.**
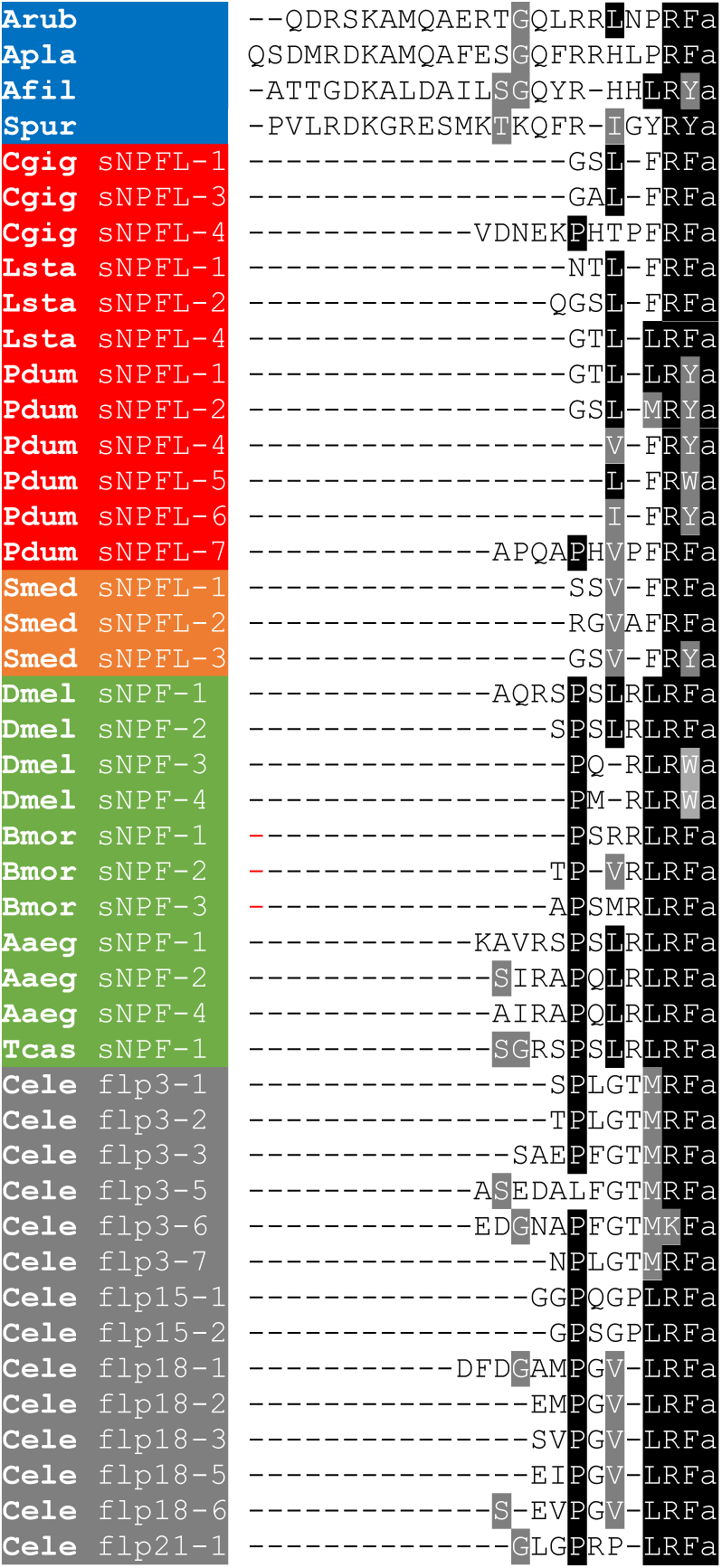
Comparison of the sequences of Ar-sNPF and orthologs from other echinoderms with protostome sNPF-type peptides. Conserved residues are highlighted in black or grey. Species names are highlighted in phylum-specific or superphylum-specific colours: blue (Echinodermata), red (Lophotrochozoa), orange-red (Platyhelminthes), green (Arthropoda) and grey (Nematoda). Species names are as follows: Aaeg (*Aedes aegypti*), Afil (*Amphiura filiformis*), Apla (*Acanthaster planci*), Arub (*Asterias rubens*), Bmor (*Bombyx mori*), Cele (*Caenorhabditis elegans*), Cgig (*Crassostrea gigas*), Dmel (*Drosophila melanogaster*), Lsta (*Lymnaea stagnalis*), Oara (*Ophiopsila aranea*), Pdum (*Platynereis dumerilii*), Smed (*Schmidtea mediterranea*), Spur (*Strongylocentrotus purpuratus*), Tcas (*Tribolium castaneum*). The accession numbers of the sequences included in this alignment are listed in supplementary table 4.

### Comparison of the structure of genes encoding precursors of sNPF-type peptides

Having identified the 22-residue amidated peptide pQDRSKAMQAERTGQL<u>RR</u>LNPRF-NH_2_ (Ar-sNPF) as the ligand for Ar-sNPFR, it was also of interest to compare the structure of genes encoding orthologs of this peptide in echinoderms for which genome sequence data are available with the structure of genes encoding sNPF-type peptides in protostomes (Figure 6). Consistent with the variability in the number of neuropeptides derived from sNPF-type precursors, we found that the structure of the genes encoding these proteins was also highly variable. Thus, the number of introns interrupting the coding sequence ranges from one in the starfish *A. planci* and in the mollusc *C. gigas* to as many as five in the *C. elegans* FLP-15 precursor gene. However, a consistent feature is the presence of an intron located after the protein-coding exon(s) that encode the N-terminal signal peptide. It is noteworthy that in the echinoderm precursor genes this intron interrupts the coding sequence for the sNPF-type peptide, whereas in protostome sNPF-type genes the coding sequences for sNPF-type peptides are located 3’ to this intron. This intron may be an evolutionarily conserved feature of sNPF-type precursor genes in the Bilateria, but with there being a shift in the position of the cleavage site that precedes the sNPF-type neuropeptide in echinoderm precursor proteins. A shift N-terminally in the location of the cleavage site would also explain why echinoderm sNPF-type peptides are longer than protostome sNPF-type peptides. Alternatively, the structure of genes encoding sNPF-type precursors in echinoderms might represent the ancestral condition in the Bilateria, and the occurrence of shorter sNPF-type peptides in protostomes could be explained by a C-terminally directed shift in the cleavage site that precedes the first or only neuropeptide derived from sNPF-type precursors.

**Figure 6.**
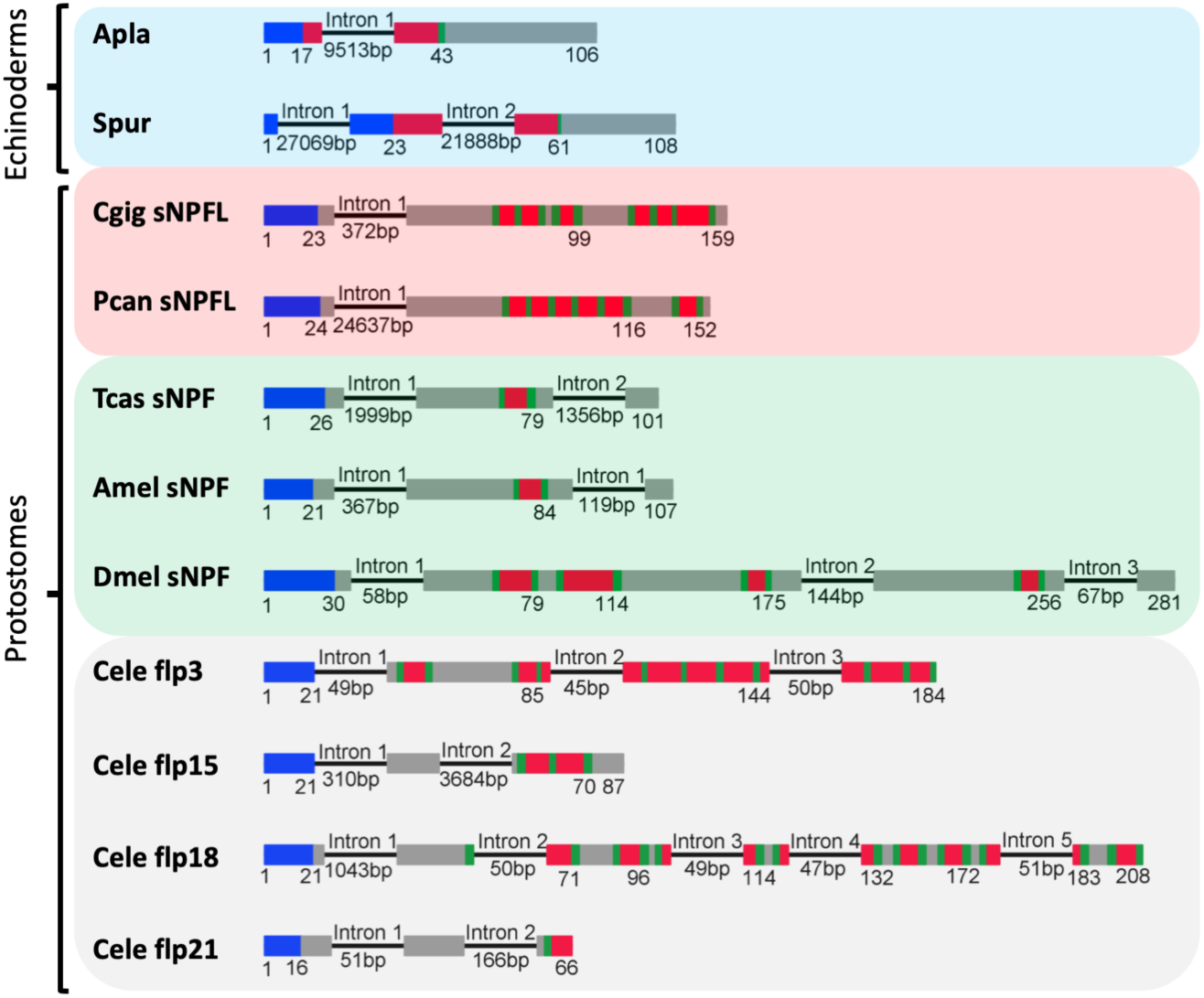
Comparison of the exon/intron structure of genes encoding echinoderm orthologs of the Ar-sNPF precursor and genes encoding protostome sNPF-type precursors. Schematic representations of the gene structures are shown, with protein-coding exons shown as rectangles and introns shown as lines (with intron length stated underneath). The protein-coding exons are colour-coded to show regions that encode the N-terminal signal peptide (blue), the neuropeptide(s) (red), monobasic or dibasic cleavage sites (green) and other regions of the precursor protein (grey). The coloured backgrounds label the following the taxonomic groups: echinoderms (blue), lophotrochozoa (red), arthropods (green) and nematodes (grey). Species abbreviations: Apla (*Acanthaster planci*), Spur (*Strongylocentrotus purpuratus*), Cgig (*Crassostrea gigas*), Pcan (*Pomacea canaliculata*), Tcas (*Tribolium castaneum*), Amel (*Apis mellifera*), Dmel (*Drosophila melanogaster*), Cele (*Caenorhabditis elegans*). The accession numbers for the sequences of the precursors shown in this figure are listed in supplementary table 5.

### Comparison of the sequences of echinoderm sNPF-type peptides with related peptides in other invertebrate deuterostomes and with vertebrate prolactin-releasing peptides

Based on a cluster analysis of neuropeptide receptor relationships, it has been proposed previously that protostome sNPF-type signalling may be orthologous to vertebrate prolactin-releasing peptide (PrRP)-type signalling (Jékely 2013). With our discovery of sNPF-type precursors and peptides in echinoderms, a new opportunity to investigate this proposed relationship was provided. Thus, it is noteworthy that echinoderm sNPF-type peptides (22-25 residues) are similar in length to vertebrate PrRPs, which are 20-31 residues as full-length peptides and in some species can occur as N-terminally truncated peptides due the presence of a monobasic cleavage site (Hinuma et al. 1998; Tachibana and Sakamoto 2014). Furthermore, by analysing sequence data from the hemichordate *S. kowalevskii* and the cephalochordate *B. floridae* here we identified novel neuropeptides that share sequence similarity with echinoderm sNPF-type peptides and with vertebrate PrRPs (Figure 7A). Thus, sequence alignment reveals that, in addition to a shared characteristic of a C-terminal RFamide or a RYamide (Y and F being synonymous substitutions), there are thirteen other residues in chordate PrRPs that are identical or structurally similar to equivalently positioned residues in at least one of the echinoderm sNPF-type peptides or the hemichordate PrRP-like peptides, as highlighted by the asterisks in Figure 7A. Thus, the discovery of echinoderm sNPF-type peptides and related peptides in other invertebrate deuterostomes has provided important new evidence that is supportive of the hypothesis that protostome sNPF-type neuropeptides and vertebrate PrRPs are orthologous.

**Figure 7.**
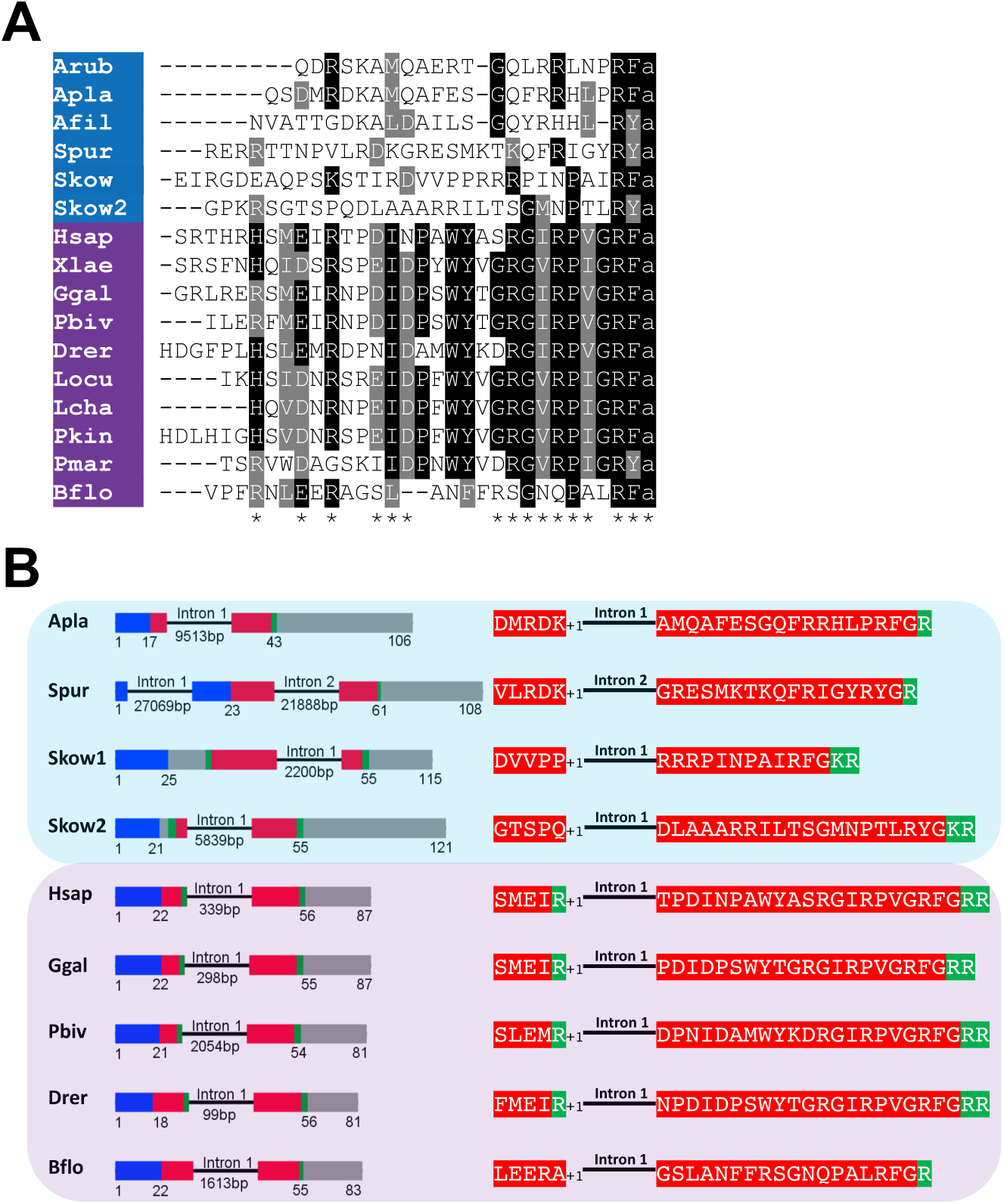
Comparison of the sequences and gene structure of echinoderm sNPF peptides, chordate prolactin-releasing peptides (PrRP) and PrRP-like peptides in the hemichordate *S. kowalevskii*. A) Sequence alignment of echinoderm sNPF-type peptides with chordate PrRP-type peptides and PrRP-like peptides in the hemichordate *S. kowalevskii*. Conserved residues are highlighted in black or grey. The asterisks indicate residues that are conserved between chordate (purple) PrRP-type peptides and at least one of the ambulacrarian (blue) peptides. B) Comparison of exon/intron structure of genes encoding echinoderm precursors of sNPF-type peptides, hemichordate precursors of PrRP-like peptides and chordate PrRP-type peptides. Schematic representations of the gene structures are shown, with protein-coding exons shown as rectangles and introns shown as lines (with intron length stated underneath). The protein-coding exons are colour-coded to show regions that encode the N-terminal signal peptide (blue), the neuropeptide(s) (red), monobasic or dibasic cleavage sites (green) and other regions of the precursor protein (grey). Note that a common characteristic is that an intron interrupts the coding sequence in the N-terminal or central region of the neuropeptide, with the intron consistently located between the first and second nucleotides (represented by the +1) of the codon for the amino acid shown after intron. Species are highlighted in clade-specific colours: blue (Ambulacraria), purple (Chordata). Species names are as follows: Arub (*Asterias rubens*), Apla (*Acanthaster planci*), Afil (*Amphiura filiformis*), Spur (*Strongylocentrotus purpuratus*), Skow, (*Saccoglossus kowalevskii*), Hsap (*Homo sapiens*), Xlae (Xenopus laevis), Ggal (*Gallus gallus*), Pbiv (Python bivittatus), Drer (*Danio rerio*), Lcha (*Latimeria chalumnae*), Locu (*Lepisosteus oculatus*), Pkin (*Paramormyrops kingsleyae*), Pmar (*Petromyzon marinus*), Bflo (*Branchiostoma floridae*). The accession numbers of the sequences included in this alignment are listed in supplementary table 6.

### Comparison of the structure of genes encoding echinoderm sNPF-type peptides, vertebrate prolactin-releasing peptides and related peptides in other invertebrate deuterostomes

Having found that echinoderm sNPF-type peptides share sequence similarity with vertebrate PrRPs, we also compared the structure of genes encoding the precursors of these peptides (Figure 7B). This revealed that a common characteristic is the presence of an intron that interrupts the coding sequence at a position corresponding to the N-terminal or central region of the echinoderm sNPFs and vertebrate PrRPs. Furthermore, in both echinoderm sNPF-type genes and vertebrate PrRP genes the intron interrupts the coding sequence in the same frame, at a position between the first and second nucleotide of the interrupted codon, which is denoted by +1 in Figure 7B. Genes encoding novel precursors of PrRP-like peptides in *S. kowalevskii* and *B. floridae* also have an intron in the +1 frame. Furthermore, in the *B. floridae* gene and in one of the *S. kowalevskii* genes (Skow 2) the intron is located in the region of the gene encoding the N-terminal part of the neuropeptide, whereas in the other *S. kowalevskii* gene (Skow1) the intron is located in a region encoding the C-terminal part of the neuropeptide. The presence of a conserved intron in the same frame in echinoderm sNPF-type genes, the two *S. kowalevskii* PrRP-like neuropeptide precursor genes and chordate PrRP-type genes supports the hypothesis that echinoderm sNPF-type neuropeptides are orthologs of hemichordate PrRP-like and chordate PrRP-type neuropeptides.

## Discussion

### NPY-like peptides in echinoderms are not orthologs of NPY/NPF-type neuropeptides

Precursors of peptides that share sequence similarity with members of the bilaterian NPY/NPF-type neuropeptide family were discovered recently in the phylum Echinodermata (Zandawala et al. 2017). These include proteins identified in several brittle star species (class Ophiuroidea) and the starfish species *Asterias rubens* and *Patiria miniata* (class Asteroidea). Here we report the cloning and sequencing of a cDNA encoding the precursor of the NPY-like peptide in *A. rubens*. Furthermore, the primary structure of this peptide (ArNPYLP) was determined using mass spectrometry, demonstrating that it is a twenty two-residue amidated peptide with an N-terminal pyroglutamate. However, comparison of the sequences of ArNPYLP and related peptides from other echinoderms with NPY/NPF-type neuropeptides from other bilaterians revealed some striking differences. Most notable is that the echinoderm peptides lack two conserved proline residues that are present in the majority of NPY/NPF-type neuropeptides that have been identified in other bilaterians. These prolines form part of what is known as the polyproline-helix or polyproline-fold, which interacts with other conserved residues (a leucine residue and two tyrosine residues) that have been shown to be important in determining the three-dimensional structure of NPY-type peptides in vertebrates (Blundell et al. 1981; Glover et al. 1983; Glover et al. 1984). Furthermore, it has been shown that these residues in human NPY are important for receptor activation (Schwartz et al. 1990; Keire et al. 2000; Nygaard et al. 2006). Although there have been no structural studies on NPF-type peptides from protostomes, our alignment (Figure 1) reveals that residues important in determining the secondary structure of vertebrate NPYs are conserved in protostome NPFs. For example, the Y/F residue in position 24, the I/L residue in position 28 and the Y residue in position 31 are conserved in both NPF-type and NPY-type peptides. These residues have been shown to be important for the formation of the three-dimensional structure in vertebrate NPY-type peptides (Blundell et al. 1981; Glover et al. 1983; Glover et al. 1984) so these residues may likewise be important for NPF receptor activation and NPF bioactivity. Furthermore, these conserved residues are also present in an NPY/NPF-type peptide that has been identified in *S. kowalevskii*, a species belonging to the phylum Hemichordata, which is a sister phylum to the Echinodermata in the ambulacrarian clade of the Bilateria (Mirabeau and Joly 2013; Elphick and Mirabeau 2014). The absence of these conserved residues in the NPY/NPF-like peptides that have been identified in *A. rubens* and other echinoderms suggests, therefore, that these peptides are not orthologs of NPY-type neuropeptides.

Our analysis of the structure of genes encoding the echinoderm NPY-like peptides provided further evidence that these peptides are not orthologs of NPY-type neuropeptides. Previous studies have compared the structure of genes encoding vertebrate NPY-type peptides and invertebrate NPF-type peptides (Blomqvist et al. 1992; Mair et al. 2000). Specifically, comparison of the structure of the human NPY precursor gene with the structure of the gene encoding the NPF precursor in the platyhelminth *Moniezia expansa* revealed that in both genes the first protein-coding exon encodes the N-terminal signal peptide and most of the NPY/NPF-type peptide through to the first two nucleotides of the codon for the penultimate residue, an arginine residue. The next exon contains the third nucleotide of the arginine codon and codons for i) a C-terminal tyrosine (in the case of the human NPY gene) or a C-terminal phenylalanine (in the case of *M. expansa* NPF gene), ii). a glycine residue that is a substrate for C-terminal amidation, iii). a dibasic cleavage site (KR) and iv). part of the C-terminal region of the precursor protein (Mair et al. 2000). Here we expanded comparative analysis of NPY/NPF gene structure to include other bilaterians. We found that a gene structure in which most of NPY/NPF-type neuropeptide sequence is encoded in one exon and the C-terminal F or Y and the amidation and cleavage sites are in the next exon is a highly conserved feature of NPY/NPF genes, which is seen in vertebrates, cephalochordates, hemichordates, lophotrocozoans, priapulids, arthropods and nematodes. Therefore, our finding that this is not a feature of genes encoding the NPY/NPF-like peptides in echinoderms (the starfish *A. planci* and the sea urchin *S. purpuratus*) provides important further evidence that these peptides are not orthologs of NPY/NPF-type neuropeptides.

### Discovery of sNPF-type neuropeptide signalling in echinoderms

If the echinoderm NPY/NPF-like peptides are not orthologs of the NPY/NPF-type neuropeptide family, then a logical prediction would be that orthologs of receptors for NPY/NPF-type neuropeptides are also absent in echinoderms, because analysis of neuropeptide-receptor co-evolution in the Bilateria has revealed that loss of a neuropeptide in an animal lineage is invariably accompanied by loss of its cognate receptor (Mirabeau and Joly 2013). Therefore, we performed a detailed phylogenetic analysis of sequence data to address this issue. Consistent with our prediction, orthologs of bilaterian NPY/NPF-type receptors were not found in any of the echinoderm species analysed. However, we discovered that *A. rubens* and other echinoderms do have orthologs of sNPF-type receptors, paralogs of the NPY-type receptors that hitherto have only been characterised in protostomes. Therefore, we hypothesised that the echinoderm NPY-like peptides may act as ligands for sNPF-type receptors and performed experimental studies to test this hypothesis. Having identified a transcript encoding a sNPF-type receptor in the starfish *A. rubens* (Ar-sNPFR), we cloned a cDNA encoding this receptor and expressed it in CHO-K1 cells. Then the *A. rubens* NPY-like peptide (ArNPYLP) was tested as a candidate ligand for Ar-sNPFR. This revealed that ArNPYLP causes dose-dependent activation of the Ar-sNPFR with an EC_50_ value of 0.15 nM, demonstrating that it is a potent ligand for this receptor. Evidence of the specificity of peptide-receptor pairing was established by our finding that other peptides, including a C-terminal fragment of ArNPYLP (LNPRFamide) and the *A. rubens* luqin-type neuropeptide ArLQ (Yañez-Guerra et al. 2018), do not act as ligands for Ar-sNPFR. Therefore, we conclude that the twenty two-residue neuropeptide formerly referred to as ArNPYLP is the natural ligand for the *A. rubens* sNPF-type receptor Ar-sNPFR and therefore this peptide should be renamed Ar-sNPF. Our discovery of the Ar-sNPF – Ar-sNPFR signalling system in *A. rubens* is important because this is the first sNPF-type signalling system to be identified in a deuterostome. Thus, sNPF-type signalling is not unique to protostomes, as has been suggested previously, and the evolutionary origin of this signalling system can be traced back to the common ancestor of the Bilateria.

Our discovery of sNPF-type signalling in a deuterostome, the starfish *A. rubens* (Phylum Echinodermata) and our and previous (Mirabeau and Joly 2013) phylogenetic analyses of neuropeptide receptor relationships indicates that sNPF-type and NPY/NPF-type signalling are paralogous. Thus, we can infer that gene duplication in a common ancestor of the Bilateria gave rise to paralogous NPY/NPF-type and sNPF-type precursor genes and paralogous NPY/NPF-type and sNPF-type receptor genes. In this context, by analysing the phylogenetic distribution and sequences of NPY/NPF-type and sNPF-type precursors and receptors, the evolutionary history of these signalling systems in the Bilateria can be examined and reconstructed.

### Reconstructing the evolution of NPY/NPF-type neuropeptide signalling

Our analysis and previous analysis (Mirabeau and Joly 2013) of the phylogenetic distribution of NPY/NPF-type signalling indicates that this neuropeptide system has been widely preserved and is highly conserved in the Bilateria, with relatively few instances of loss based on the data currently available (Figure 8). Thus, genes encoding NPY/NPF-type precursors and genes encoding proven or candidate receptors for NPY/NPF-type peptides have been identified in deuterostomes (vertebrates, hemichordates) and protostomes (platyhelminthes, annelids, molluscs, arthropods). Furthermore, here we report the first identification of genes encoding a NPY/NPF-type precursor and a NPY/NPF-type receptor in the protostome phylum Priapulida. It is noteworthy that NPY/NPF-type signalling has only been partially characterised in a model invertebrate system – the nematode *C. elegans*. Our phylogenetic analysis indicates that there are two *C. elegans* receptors that are orthologs of NPY/NPF-type receptors: NPR-12, which is an orphan receptor, and NPR-11, which has been shown to be activated by the peptide MDANAFRMSFamide (Chalasani et al. 2010). However, this peptide shares little sequence similarity with NPY/NPF-type peptides from other bilaterians. Furthermore, receptor assays only showed activation at peptide concentrations of 10 and 30 µM (Chalasani et al. 2010), which are high when compared to other reported NPY/NPF-type receptors that are typically activated in the nanomolar range (Bard et al. 1995; Lundell et al. 1997; Garczynski et al. 2002; Saberi et al. 2016). Recently, based on similarity-based sequence alignments, it has been suggested that the mature peptide derived from the *C. elegans* protein FLP-27 may be an ortholog of NPY/NPF-type peptides (Fadda et al. 2019). Here, our analysis of the structure of the gene encoding the FLP-27 precursor has revealed that it has the characteristic structure of NPY/NPF-type genes, with an intron interrupting the codon for the C-terminal arginine of the NPF-type peptide sequence. Thus, based on our analysis of *C. elegans* sequence data, we conclude that the NPY/NPF-type peptide derived from the FLP-27 precursor protein is likely to act as a ligand for the NPR-11 and/or NPR-12 receptors.

**Figure 8.**
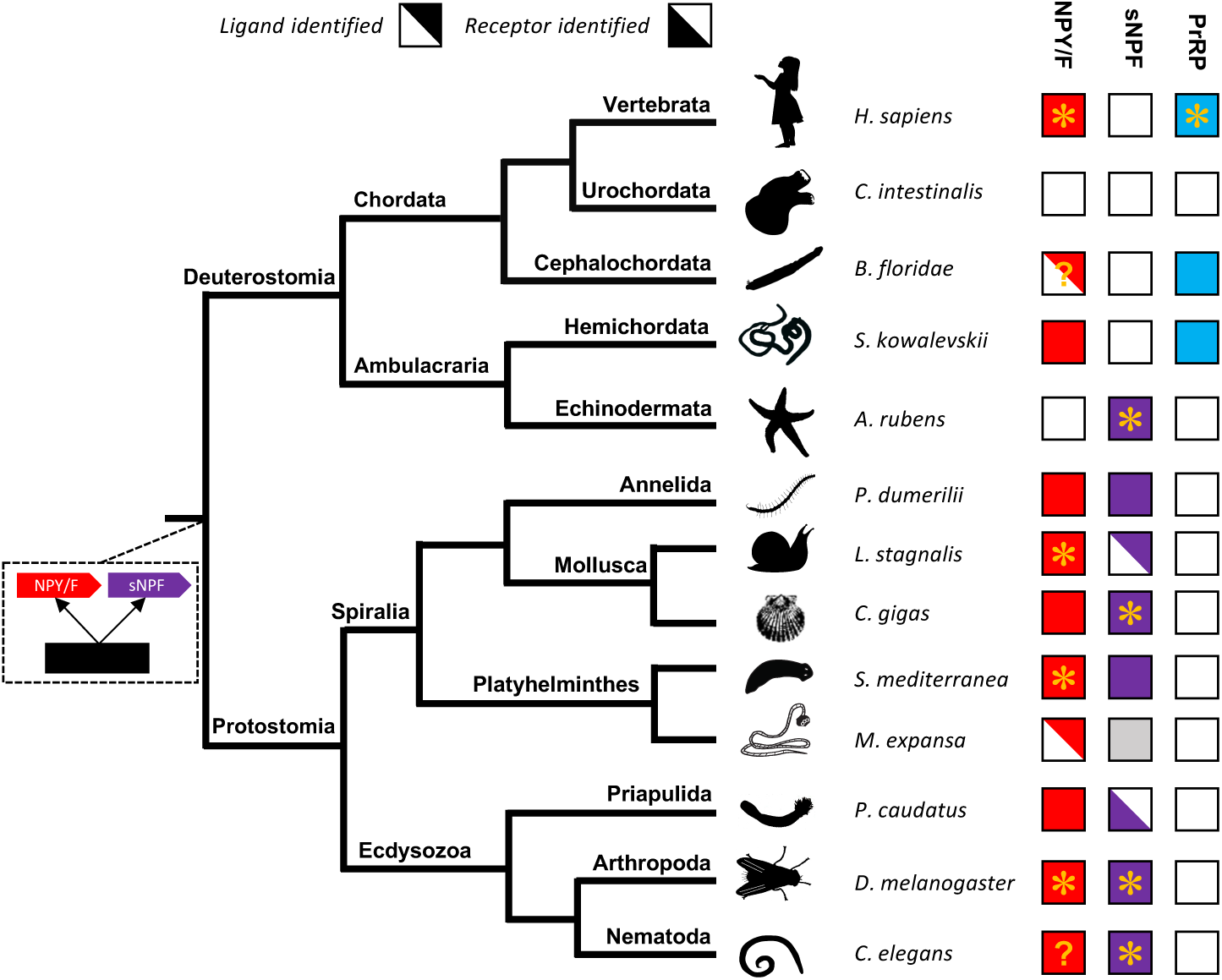
Phylogenetic diagram showing the occurrence of NPY/NPF-type, sNPF-type and PrRP-type neuropeptide signalling in the Bilateria. The phylogenetic tree shows relationships of selected bilaterian phyla. Phyla in which NPY/NPF-type peptides/precursors and NPY/NPF receptors have been identified are labelled with red-filled squares. Phyla in which sNPF-type peptides/precursors and sNPF-type receptors have been identified are labelled with purple-filled squares. Phyla in which PrRP-type peptides/precursors and PrRP-type receptors have been identified are labelled with blue-filled squares. The inclusion of an asterisk in filled squares indicates that activation of a receptor by a peptide ligand has been demonstrated experimentally. Note that the starfish *Asterias rubens* is the first and only deuterostome in which the neuropeptide ligand for an sNPF-type receptor has been identified. Note also the mutually exclusive patterns in the phylogenetic distribution of sNPF-type signalling and PrRP-type signalling, with the former found only in protostomes and echinoderms and the latter found only in vertebrates, cephalochordates and hemichordates, which is supportive of the hypothesis that these signalling systems may be orthologous. NPY/NPF-type signalling occurs in most phyla, but it has been lost in echinoderms and urochordates. However, the inclusion of a question mark for the putative NPY/NPF-type peptide identified in the cephalochordate *B. floridae* (Mirabeau and Joly 2013; Elphick and Mirabeau 2014) signifies that it is atypical of NPY/NPF-type peptides, which may explain why NPY/NPF-type receptors have yet to be identified in cephalochordates. The inclusion of a question mark in the *C. elegans* red square indicates that the peptide identified as a ligand for the *C. elegans* NPY/NPF-type receptor (Chalasani et al. 2010) does not have the typical features of an NPY/NPF-type peptide. The grey square for sNPF in *M. expansa*, for which only transcriptome sequence data are available, indicates that sNPF-type peptides and sNPF-type receptor(s) are likely to be present in this species because sNPF-type peptides and sNPF-type receptors have been identified in another platyhelminth species, *S. mediterranea*, for which a genome sequence is available. Species names are as follows: *H. sapiens (Homo sapiens), C. intestinalis (Ciona intestinalis), B. floridae (Branchiostoma floridae), S. kowalevskii (Saccoglossus kowalevskii), A. rubens (Asterias rubens), P. dumerilii (Platynereis dumerilii), L. stagnalis (Lymnaea stagnalis), M. expansa* (*Moniezia expansa*), *S. mediterranea* (*Schmidtea mediterranea*), *C. gigas (Crassostrea gigas), D. melanogaster (Drosophila melanogaster), C. elegans (Caenorhabditis elegans).* Silhouettes of representative animals from each phylum are from www.openclipart.com and they are free from copyright.

The first NPF-type peptide was discovered in the platyhelminth *Monieza expansa* (Maule et al. 1991), but the receptor for this ligand has not been identified. We were unable to identify a candidate receptor for NPF in *M. expansa,* which probably reflects the limited availability of sequence data for this species. However, genome/transcriptome sequence data are available for the flatworm species *S. mediterranea* and an expanded family of sixteen putative NPY/NPF-type receptors (*Smed*-NPYR1 - *Smed*-NPYR16) in this species has been reported (Saberi et al. 2016). Conversely, our phylogenetic analysis (Figure 2) has revealed that only *Smed*-NPYR1, *Smed*-NPYR3, *Smed*-NPYR5 and *Smed*-NPYR6, are orthologs of the NPY/NPF-type family of receptors. Accordingly, it has been shown that Smed-NPYR1 is activated by a peptide that has a characteristic NPY/NPF-type structure (Saberi et al. 2016).

In the annelid *P. dumerilii*, a receptor named NPY-4 receptor 1 that is activated by three NPY/NPF-type peptides (NPY1, NPY3 and NPY4) has been reported previously. However, cluster analysis indicated that this receptor may not be an NPY/NPF-type receptor (Bauknecht and Jékely 2015). Interestingly, we have identified an NPF-type precursor and peptide in *P. dumerilii* that has not been reported previously and which contains residues that are conserved in NPF-type peptides from other protostomes (Figure 1). Furthermore, the exon/intron structure of a gene encoding an ortholog of the *P. dumerilii* NPF-type precursor in the annelid *Helobdella robusta* is consistent with NPY/NPF-type precursor genes (Figure 2). It is likely, therefore, that the NPF-type peptide in *P. dumerilii* is the ligand for the orphan receptor GPR62, which is clearly an ortholog of NPY/NPF-type receptors (Figure 3).

Although NPY/NPF-type signalling has been retained in the majority of phyla, as discussed above, it has been reported previously that NPY/NPF-type signalling has been lost in urochordates (Mirabeau and Joly 2013). Furthermore, here we present evidence for the first time indicating that NPY/NPF-type signalling has also been lost in echinoderms. The functional significance of the loss of NPY/NPF-type signalling in urochordates and echinoderms is unknown. However, insights into this issue may emerge as we learn more about the physiological roles of NPY/NPF-type signalling in a variety of invertebrate taxa.

### Reconstructing the evolution of sNPF-type neuropeptide signalling

Discovery of sNPF-type neuropeptide signalling in echinoderms is interesting because orthologs of protostome sNPF-type receptors have not been identified in other deuterostome phyla – Chordata and Hemichordata. Conversely, both peptides and receptors of the sNPF-type signalling system have been identified in several protostome phyla (Figure 8). In this context, it is of interest to first review here what is currently known about the molecular components of the sNPF-type signalling system in protostomes.

Starting with the ecdysozoan protostomes, it was originally thought that sNPF-type signalling may be arthropod-specific, reflecting the original discovery of this signalling system in insects (Nässel and Wegener 2011). However, a large-scale phylogenetic analysis of G-protein coupled neuropeptide receptors in the Bilateria revealed an expanded family of genes encoding sNPF-type receptors in the nematode *C. elegans* (Mirabeau and Joly 2013). Our phylogenetic analysis confirms the existence of an expanded family of sNPF-type receptors in *C. elegans* (Figure 2). Furthermore, we show that the *C. elegans* receptors NPR1, NPR2, NPR3, NPR4 and NPR5, which are activated by sNPF-type peptides derived from the FLP-15, FLP-18 and FLP-21 precursors (Kubiak, Larsen, Nulf, et al. 2003; Kubiak, Larsen, Zantello, et al. 2003; Rogers et al. 2003; Kubiak et al. 2008; Cohen et al. 2009; Ezcurra et al. 2016), form part of a clade of sNPF-type receptors, together with deorphanised sNPF-type receptors from the insects *D. melanogaster* (Mertens et al. 2002), *B. mori* (Yamanaka et al. 2008; Ma et al. 2017) and *A. aegypti* (Christ et al. 2018). Previously, NPR1, NPR2 and NPR5 were annotated as NPY/NPF-type receptors and NPR3 and NPR4 were annotated as NPY/NPF-like receptors (Cardoso et al. 2012) but it is now clear that NPR1-5 are in fact sNPF-type receptors. Hitherto the existence of sNPF-type signalling in priapulids has not been reported. Here our analysis of sequence data from *Priapulus caudatus* has identified an sNPF-type receptor but we did not identify a precursor protein that gives rise to a candidate ligand for this receptor and therefore this is an objective for future work.

Turning to the spiralian protostomes, sNPF-type receptors were identified in molluscs, annelids and platyhelminths (Figure 2, 8). However, the peptide ligand(s) for sNPF-type receptors have only been demonstrated experimentally in a single molluscan species – the bivalve *C. gigas* (Bigot et al. 2014), although orthologs of these peptides were functionally characterised in other molluscan species prior to this (Hoek et al. 2005; Zatylny-Gaudin et al. 2010; Zhang et al. 2012). Interestingly our phylogenetic analysis (Figure 2) revealed that a clade comprising spiralian molluscan sNPF-type receptors also contains a receptor from the annelid *Platynereis dumerilli* that has been experimentally characterised as a receptor that is activated by the amidated tetrapeptide FMRFamide and a peptide known as NKY (Bauknecht and Jékely 2015). Furthermore, the NKY peptide and the NKY receptors have been described as paralogs of NPY-type peptides and NPY-type receptors, respectively (Bauknecht and Jékely 2015). Although our phylogenetic analysis indicates that the *C. gigas* sNPF receptor and the *P. dumerilli* NKY receptor are orthologs, there is a discrepancy in the ligands that activate these two receptors. The *C. gigas* sNPF-type receptor is activated by a sNPF-type peptide comprising five to six residues and with a C-terminal LFRFamide sequence (Bigot et al. 2014), whereas the *P. dumerilii* NKY receptor was shown to be activated by NKY-type peptides that are typically up to forty-three residues in length and with a C-terminal LLRYamide sequence (Bauknecht and Jékely 2015). Therefore, although it was not the primary purpose of this study, we investigated this anomaly by comparing the ability of three peptides to act as ligands for the *C. gigas* sNPF receptor: i). the peptide GSLFRFamide, which has been shown previously to act as a ligand for this receptor (Bigot et al. 2014), ii). the amidated tetrapeptide FMRFamide and iii). a *C. gigas* NKY-type peptide. This experiment revealed that GSLFRFamide is the most potent ligand of this receptor, with an EC_50_ value of 31 nM (Supplementary Figure 5). Interestingly, however, we found that the *C. gigas* NKY-type peptide and FMRFamide also cause activation of the receptor, but only at relatively high concentrations. Thus, the EC_50_ for FMRFamide was 3.4 µM and the EC_50_ for the *C. gigas* NKY-type peptide was 3.02 µM. We conclude from this that GSLFRFamide is a natural ligand for the *C. gigas* sNPF receptor, consistent with the findings of (Bigot et al. 2014), whereas the ability of FMRFamide and the *C. gigas* NKY-type peptide to activate the *C. gigas* sNPF-type receptor may reflect non-physiological neuropeptide-receptor cross-talk. Accordingly, the *P. dumerilii* receptor identified as a receptor for NKY (Bauknecht and Jékely 2015) may be activated physiologically by shorter sNPF-type GSLFRFamide-like peptides, the sequences of which we show in the alignment in Figure 5 (e.g. GTLLRYamide, GSLMRYamide etc.). It is noteworthy that the C-terminal tetrapeptide of GTLLRYamide is similar to the C-terminal tetrapeptide of *P. dumerilli* NKY-1 (IMRYamide), which likely explains why NKY was found to act as a ligand, albeit with an EC_50_ of 420 nM, for a *P. dumerilli* NKY/sNPF receptor (Bauknecht and Jékely 2015). Further studies are now needed to investigate the ligand-binding properties of the *P. dumerilli* NKY/sNPF receptor in more detail.

As highlighted above, an expanded family of sixteen putative NPY/NPF-type receptors (*Smed*-NPYR1 - *Smed*-NPYR16) has been identified in the platyhelminth *S. mediterranea* (Saberi et al. 2016). However, our phylogenetic analysis indicates that four of these receptors (*Smed*-NPYR7, *Smed*-NPYR8, *Smed*-NPYR9, and *Smed*-NPYR10) are orthologs of sNPF-type receptors. Therefore, it would be expected that the peptide ligands for these receptors are similar to the peptides that have been identified as ligands for sNPF-type receptors in another spiralian – the mollusc *C. gigas* (Bigot et al. 2014). On this basis, we have identified candidate ligands for *S. mediterranea* sNPF-type receptors, which are included in the alignment shown in Figure 5 (SSVFRFamide, RGVAFRFamide and GSVFRYamide).

Having reviewed the characteristics of sNPF-type signalling in protostomes, it is of interest to make comparisons with the sNPF-type signalling system that has been identified here for the first time in a deuterostome phylum – the Echinodermata. Alignment of the sequences of protostome sNPF-type peptides with the echinoderm sNPF-type peptides reveals modest C-terminal sequence similarity, as shown in Figure 5 and as described in the results section of this paper. Furthermore, the echinoderm sNPF-type peptides are much longer than protostome sNPF-type peptides. Another difference is that protostome sNPF-type neuropeptide precursors typically give rise to multiple sNPF-type peptides, whereas in echinoderms the sNPF-type precursor contains a single sNPF-type peptide that is located adjacent to the signal peptide. Likewise, comparison of the structure of the genes encoding sNPF-type precursors in protostomes and echinoderms reveals limited similarity (Figure 6). Thus, there is little evidence of orthology from comparison of the neuropeptide, precursor and gene sequences in protostomes and echinoderms. Consequently, our conclusion that the echinoderm NPY-like peptides are orthologs of protostome sNPF-type peptides is principally based on the orthology of their receptors, as shown in Figure 3. It is important to note, however, that this is not unprecedented in investigations of the evolution neuropeptide signalling. Thus, whilst the sequences of some neuropeptides and neuropeptide precursors are highly conserved throughout the Bilateria, others are so divergent that they can be unrecognisable as orthologs. An example of the former are vasopressin/oxytocin (VP/OT)-type neuropeptides and precursors. An example of the latter are neuropeptide-S (NPS)/crustacean cardioactive peptide (CCAP)-type neuropeptides and precursors, which are paralogs of VP/OT-type neuropeptides and precursors (Semmens et al. 2015). Thus, by way of comparison, NPY/NPF-type neuropeptides are similar to VP/OT-type neuropeptides in exhibiting a high level of sequence conservation throughout the Bilateria. Conversely, sNPF-type neuropeptides are similar to NPS/CCAP-type neuropeptides in being highly divergent, with neuropeptides in protostomes and deuterostomes exhibiting modest sequence similarity.

The discovery of sNPF-type signalling in echinoderms has provided a unique opportunity to speculate on the ancestral characteristics of this signalling system in Urbilateria. It is noteworthy that, by comparison with the protostome sNPF-type peptides, the echinoderm sNPF-type peptides have more features in common with the paralogous NPY/NPF-type peptides. The echinoderm sNPF-type peptides are not as long as NPY/NPF-type peptides but they are nevertheless much longer than protostome sNPF-type peptides. Furthermore, it was the sequence similarity that echinoderm peptides share with NPY/NPF-type peptides that originally facilitated their discovery (Zandawala et al., 2017). Additionally, the structure of the echinoderm sNPF-type precursors is similar to NPY/NPF-type precursors because the neuropeptide is located immediately after the signal peptide, whereas this is not a feature of protostome sNPF-type precursors. Based on these observations, we propose that echinoderm sNPF-type peptides and precursors may more closely resemble the ancestral characteristics of this signalling system in Urbilateria. Furthermore, we speculate that the common ancestor of the paralogous NPY/NPF-type and sNPF-type neuropeptide precursors may have been similar to NPY/NPF-type precursors with respect peptide, precursor and gene structure. Then, following gene duplication, these ancestral characteristics were retained in the paralog that gave rise to the bilaterian NPY/NPF-type peptides/precursors. In contrast, the paralog that gave rise to sNPF-type signalling diverged from the ancestral condition. However, the extent of divergence varies in the echinoderm and protostome lineages. In echinoderms, the sNPF-type peptides/precursors have many NPY/NPF-type characteristics and we conclude that this reflects less divergence from the proposed ancestral condition. Conversely, in the protostomes, the sNPF-type peptides/precursors exhibit little similarity with NPY/NPF-type peptides/precursors and we conclude that this reflects more divergence from the proposed ancestral condition.

Lastly, we need to consider more broadly the evolutionary history of sNPF-type signalling in deuterostomes, and in particular the non-echinoderm phyla – the hemichordates and chordates. A detailed phylogenetic analysis of G-protein coupled neuropeptide receptors in the Bilateria did not reveal the presence of orthologs of protostome sNPF-type receptors in hemichordates and chordates (Mirabeau and Joly 2013) and likewise our more specific analysis of the phylogenetic distribution and relationships NPY/NPF-type and sNPF-type receptors (Figure 3) also did not reveal the presence of orthologs of protostome sNPF-type receptors in hemichordates and chordates. Based on these findings and our discovery of sNPF-type signalling in echinoderms it could be inferred that sNPF-type signalling has been lost in hemichordates and chordates. However, a cluster analysis of bilaterian neuropeptide receptor relationships has revealed that protostome sNPF-type receptors cluster with receptors for vertebrate prolactin-releasing peptides (PrRPs) (Jekely, 2013). This contrasts with our (Figure 3) and previous (Mirabeau and Joly 2013) analysis of neuropeptide receptor relationships by generation of phylogenetic trees, which revealed that chordate and hemichordate PrRP-type receptors do not clade with sNPF-type receptors. A limitation of cluster analysis of receptor relationships is that it is based on pairwise comparisons that cannot resolve paralogy/orthology relationships because speciation/duplication nodes are not retrieved (Gabaldón 2008; Kim et al. 2008). Thus, the determination of deep homology relationships is normally accomplished by generating phylogenetic trees (Gabaldón 2008). Nevertheless, informed by the hypothesis of Jekely (2013) that sNPF-type signalling may be orthologous to PrRP-type signalling, which was also reported in a recent review article (Fadda et al., 2019), here we investigated the occurrence PrRP-like neuropeptides in the hemichordate *S. kowalevskii* and the cephalochordate *B. floridae*. Importantly, we identified one precursor protein in *B. floridae* and two precursor proteins in *S. kowalevskii* that contain peptides that share sequence similarity with both echinoderm sNPF-type neuropeptides and with vertebrate PrRP-type neuropeptides. Thus, as shown in the alignment in Figure 7A, in addition to a conserved RF/RY-amide C-terminal group there are thirteen other residues in the chordate PrRP-type neuropeptides that are conserved in at least one of the echinoderm sNPF-type neuropeptides or *S. kowalevskii* PrRP-like neuropeptides. Furthermore, genes encoding echinoderm sNPFs, hemichordate PrRP-like neuropeptides and chordate PrRPs have the common characteristic of an intron that interrupts the coding sequence at a position corresponding to their N-terminal or central regions. Furthermore, the intron consistently interrupts the coding sequence in the same frame, at a position between the first and second nucleotide of the interrupted codon, as denoted by +1 in Figure 7B. This contrasts with NPY/NPF-type genes that have a highly conserved intron interrupting the coding sequence at a position corresponding to the C-terminal region of the mature peptides, with the intron located between the second and third nucleotide of the codon for a conserved arginine residue, as denoted by −1 in Figure 2. Collectively, these findings are supportive of the hypothesis that echinoderm sNPF-type neuropeptides are orthologs of vertebrate PrRP-type neuropeptides and the novel PrRP-like peptides that we have identified here in the hemichordate *S. kowalevskii* and the cephalochordate *B. floridae*. Furthermore, it is noteworthy that orthologs of vertebrate PrRP-type receptors have been identified in cephalochordates and hemichordates (Mirabeau and Joly 2013) (see also Figure 3). Thus, there are mutually exclusive patterns in the phylogenetic distribution of sNPF-type receptors and PrRP-type receptors, with the former found only in protostomes and echinoderms and the latter found only in vertebrates, cephalochordates and hemichordates (Figure 8). A parsimonious interpretation of this finding is that sNPF-type signalling is orthologous to vertebrate PrRP-type signalling. However, it is possible that duplication of an sNPF/PrRP-type signalling system occurred in a common ancestor of the deuterostomes, with paralogous signalling systems then being differentially retained in different lineages; i.e. retention of sNPF-type signalling and loss of PrRP-type signalling occuring in echinoderms and *vice versa* in hemichordates and chordates.

### General conclusions

The findings reported in this paper provide important new insights into the evolution of NPY/NPF-type and sNPF-type neuropeptide signalling systems. Discovery of a sNPF-type signalling system in an echinoderm has provided the first experimental evidence that the evolutionary origin of sNPF-type signalling can be traced back to the common ancestor of the Bilateria. Furthermore, discovery of sNPF-type neuropeptides in echinoderms has provided evidence that sNPF-type neuropeptides are orthologs of vertebrate prolactin-releasing peptides. Thus, this study powerfully illustrates the importance of research on neuropeptide signalling systems in echinoderms (and other deuterostome invertebrates) in providing key missing links for reconstruction of neuropeptide evolution.

## Material and methods

### Animals

Starfish (*Asterias rubens*) were obtained from a fisherman based at Whitstable (Kent, UK). They were then maintained in a circulating seawater aquarium at ∼11 °C in the School of Biological and Chemical Sciences at Queen Mary University of London and were fed on mussels (*Mytilus edulis*) collected near Margate (Kent, UK).

### Cloning and sequencing of a cDNA encoding the precursor of an *A. rubens* NPY-like peptide

A transcript encoding the *A*. *rubens* precursor of an NPY-like peptide (ArNPYLP) has been identified previously (GenBank: MK033631) (Zandawala et al. 2017). The cDNA containing the complete open reading frame of the ArNPYLP precursor was amplified by PCR using *A. rubens* radial nerve cord cDNA, the forward primer AAGTCAAAAGGCGAGCAAGA, the reverse primer AAAGGGATGTGGTGTTGGTG and Q5 polymerase (NEB; Cat. No. M0491S). The PCR products were ligated into the pBluescript II KS (+) vector (Invitrogen; Cat. No. K280002) that had been cut previously with the restriction enzyme *EcoRV* by performing blunt-end ligation with T4 DNA ligase (NEB; Cat. No. M0202S). The cloning was confirmed by restriction enzyme digestion and sequencing (TubeSeq service; Eurofins Genomics).

### Structural characterisation of the *A. rubens* NPY-like peptide using mass spectrometry

After confirming the nucleotide sequence of the ArNPYLP precursor by cloning and sequencing, mass spectrometry was used to determine the structure of the peptide derived from this precursor. The methods employed, including extraction of peptides from *A. rubens* radial nerve cords, treatment of samples, equilibration of columns, reverse phase chromatography for the initial separation and injection into a Orbitrap-Fusion (ThermoScientific) for tandem mass spectrometry (MS/MS), were performed using a previously reported protocol for the identification of the starfish neuropeptides (Lin et al. 2017). The methods employed for data analysis are described below. Mass spectra were searched using Sequest Proteome Discoverer (Thermo Fisher Scientific, v. 2.2) against a database comprising forty-three different precursor proteins identified by analysis of *A. rubens* neural transcriptome data, including the *A. rubens* ArNPYLP precursor and all proteins in GenBank from species belonging to the Asteriidae family and the common Repository of Adventitious Proteins Database (http://www.thegpm.org/cRAP/index.html). Theoretical peptides were generated allowing up to two missed cleavages and variable modifications, including amidation (−0.98402) of C-terminal glycines and pyroglutamate (−17.02655) of N-terminal glutamines, and oxidation of methionine (+15.99). Precursor mass tolerance was 10 ppm and fragment ions were searched at 0.8 Da tolerances. Results from Discoverer were collated and annotated in Scaffold version 4.8.4 (Proteome Software).

### Sequence alignment of the *A. rubens* NPY-like peptide with NPY-like peptides from other echinoderms and NPY/NPF-type peptides from other taxa

The amino acid sequence of ArNPYLP, as confirmed by mass spectrometry, and predicted orthologs from other echinoderm species were aligned with the sequences of NPY/NPF-type peptides from a variety of bilaterian species (see Supplementary table 1 for a list of the sequences) using MAFFT, with the number of maximum iterations set to 1000 to ensure an optimal alignment. These alignments were highlighted using the software BOXSHADE (www.ch.embnet.org/software/BOX_form.html) with 70% conservation as the minimum. Finally, the sequences were highlighted in phylum-specific or superphylum-specific colours: dark blue (Echinodermata), light blue (Hemichordata), purple (Chordata), orange (Platyhelminthes), red (Lophotrochozoa), yellow (Priapulida), green (Arthropoda), grey (Nematoda).

### Comparison of the exon/intron structure of genes encoding NPY-like peptides in echinoderms and genes encoding NPY/NPF-type peptides in other taxa

The sequences of transcripts and genes encoding precursors of echinoderm NPY-like peptides and NPY/NPF-type precursors from other taxa were obtained using BLAST (https://blast.ncbi.nlm.nih.gov/). See supplementary table 2 for a list of the transcript and gene sequences used. The online tool Splign (Kapustin et al. 2008) (https://www.ncbi.nlm.nih.gov/sutils/splign/splign.cgi) was employed to determine and analyse gene structure and schematic figures showing the gene structure were generated using IBS 1.0 (Liu et al. 2015).

### Identification and phylogenetic characterisation of an *A. rubens* G-protein coupled receptor related to NPY/NPF/sNPF-type receptors

To identify a candidate receptor for the *A. rubens* NPY-like peptide ArNPYLP, *A. rubens* neural transcriptome sequence data was analysed using sequenceserver BLAST (Priyam et al. 2015), submitting NPY-type receptors from *H. sapiens* (GenBank NP_000900.1, NP_000901.1, NP_001265724.1) an NPF-type receptor from *D. melanogaster* (GenBank AAF51909.3) and sNPF-type receptors from *D. melanogaster* (GenBank; NP_524176.1) and *C.gigas* (GenBank XP_011451552.1) as query sequences. A transcript (contig 1120879) encoding a 386-residue protein (http://web.expasy.org/translate/) was identified as the top hit in all BLAST searches and this was deposited in GenBank under the accession number MH807444. The protein sequence was also analysed using Protter V1.0 (Omasits et al. 2014). Using BLAST, homologs of the *A. rubens* protein were identified in other echinoderms, including the starfish *Acanthaster planci* (XP_022101544.1), the sea urchin *Strongylocentrotus purpuratus* (XP_003725178.1) and the sea cucumber *Apostichopus japonicus* (PIK36230.1).

To investigate the relationship of the echinoderm receptors with NPY/NPF-type receptors and sNPF-type receptors from other taxa, a phylogenetic tree was generated using the maximum-likelihood method (see supplementary table 3 for a list of sequences used). Receptor sequences were aligned using the MUSCLE plugin in MEGA 7 (iterative, 10 iterations, UPGMB as clustering method) (Edgar 2004; Kumar et al. 2016) and the alignment was manually trimmed in the C-terminal and N-terminal regions to include a total of 300 residues spanning from the first to the seventh transmembrane domains. The maximum-likelihood tree was generated using W-IQ-tree online version 1.0 (1000 bootstrap replicates, LG+G+I+F substitution model) (Trifinopoulos et al. 2016).

### Cloning and pharmacological characterisation of the *A. rubens* NPY/NPF/sNPF-type receptor

To enable the pharmacological characterisation of the *A. rubens* NPY/NPF/sNPF-type receptor, a cDNA encoding this receptor was cloned into the eukaryotic expression vector pcDNA 3.1(+) (Invitrogen; Cat. No. V790-20). To facilitate expression of the cloned receptor, the forward primer included a partial Kozak consensus sequence (ACC) and a sequence corresponding to the first 15 bases of the open reading frame of contig 1120879 (ACCATGCAGATGACAACC) and the reverse primer consisted of a stop codon and a sequence reverse complementary to the 3’ region of the open reading frame of contig 1120879 (GCGTCACATAGTGGTATCATG). PCR was performed using the forward primer and reverse primers, *A. rubens* radial nerve cord cDNA and Q5 polymerase (NEB; Cat. No. M0491S). PCR products were ligated into the pcDNA 3.1(+) vector that had been cut previously with the restriction enzyme *EcoRV* by performing blunt-end ligation with T4 DNA ligase (NEB; Cat. No. M0202S). Successful ligation and the direction of the insert was determined by restriction enzyme digestion and sequencing (TubeSeq service; Eurofins Genomics).

Chinese hamster ovary (CHO)-K1 cells stably expressing the calcium sensitive aequorin fusion protein (G5A) and transfected with the human promiscuous G-protein Gα16 (Baubet et al. 2000) were used as an expression system to functionally characterise the *A. rubens* NPY/NPF/sNPF-type receptor. Cells were cultured, transfected and luminescence assays were performed as described previously (Yañez-Guerra et al. 2018). After transfection with the *A. rubens* sNPF-type receptor, cells were exposed to the *A. rubens* NPY-like peptide pQDRSKAMQAERTGQLRRLNPRF-NH_2_ (custom synthesised by Peptide Protein Research Ltd., Fareham, UK), which was diluted in DMEM/F12 Nutrient Mixture medium at concentrations ranging from 10^−14^ M to 10^−5^ M in clear bottom 96-well plates (Sigma-Aldrich; Cat. No. CLS3603-48EA). Luminescence was measured over a 30 second period using a FLUOstar Omega Plate Reader (BMG LABTECH; FLUOstar Omega Series multi-mode microplate reader) and data were integrated over the 30-second measurement period. For each concentration, measurements were performed in triplicate, and the average of each was used to normalise the responses. The responses were normalised to the maximum luminescence measured in each experiment (100% activation) and to the background luminescence with the vehicle media (0% activation). Dose-response curves were fitted with a four-parameter curve and EC_50_ values were calculated using Prism 6 (GraphPad, La Jolla, USA), from dose–response curves based on at least three independent transfections.

### Sequence alignment of the *A. rubens* sNPF and related peptides from other echinoderms with sNPF-type peptides from other taxa

The sequences of echinoderm sNPF-type peptides were aligned with sNPF-type peptides that have been identified in protostomes (see supplementary table 4 for a list of the sequences used) using MAFFT version 7 (5 iterations, substitution matrix; BLOSUM62) and then manually curated. Highlighting of the conserved residues was done using BOXSHADE (www.ch.embnet.org/software/BOX_form.html) with 70% conservation as the minimum for highlighting.

### Comparison of the exon/intron structure of genes encoding sNPF peptides in echinoderms and genes encoding sNPF-type peptides in other taxa

The sequences of transcripts and genes encoding echinoderm sNPF-type precursors and sNPF/sNPFL/FLP-3/FLP-15/FLP-18/FLP-21 precursors from protostomes were obtained using BLAST (https://blast.ncbi.nlm.nih.gov/) (see supplementary table 5 for a list of the transcript and gene sequences analysed). The online tool Splign (Kapustin et al. 2008) (https://www.ncbi.nlm.nih.gov/sutils/splign/splign.cgi) was employed to determine the structure of genes encoding these peptide precursors. Schematic figures showing the gene structure were generated using IBS 1.0 (Liu et al. 2015).

### Sequence alignment of *A. rubens* sNPF and related peptides from other echinoderms with PrRP-type peptides from chordates and PrRP-like peptides from the hemichordate *S. kowalevskii*

To identify candidate ligands for PrRP-type receptors in the cephalochordate *B. floridae* and the hemichordate *S. kowalevskii* (Figure 3), we analysed transcriptomic and genomic sequence data for these species (Putnam et al. 2008; Simakov et al. 2015). The data analysed also included a list of predicted *S. kowalevskii* proteins kindly provided to O. Mirabeau by Dr. R.M. Freeman (Harvard Medical School, USA). The methods employed to identify candidate neuropeptide precursors have been reported previously (Mirabeau and Joly 2013) but here we had the more specific objective of identifying proteins with an N-terminal signal peptide followed by a neuropeptide with a predicted C-terminal RFamide or RYamide motif. This resulted in discovery of one candidate PrRP-type precursor in the cephalochordate *B. floridae* and two candidate PrRP-type precursors in the hemichordate *S. kowalevskii*.

The sequences of echinoderm sNPF-type peptides were aligned with chordate PrRP-type peptides and the two PrRP-like peptides from *S. kowalevskii* (see supplementary table 6 for a list of the sequences used) using MAFFT version 7 (5 iterations, substitution matrix; BLOSUM62) and then manually curated. Highlighting of the conserved residues was done using BOXSHADE (www.ch.embnet.org/software/BOX_form.html) with 70% conservation as the minimum for highlighting.

### Comparison of the exon/intron structure of genes encoding sNPF precursors in echinoderms with genes encoding precursors of PrRP-type peptides in chordates and genes encoding precursors of PrRP-like peptides in the hemichordate *S. kowalevskii*

The sequences of transcripts and genes encoding echinoderm sNPF precursors, chordate PrRP-type precursors and one of the *S. kowalevskii* precursors (Skow1) of a PrRP-like peptide were obtained using BLAST (https://blast.ncbi.nlm.nih.gov/). The sequence of a predicted transcript encoding a second *S. kowalevskii* precursor (Skow2) of a PrRP-like peptide was determined based on a GenScan prediction (Burge and Karlin 1997; Burge and Karlin 1998) from scaffold 51909 (GenBank accession number NW_003156735.1). See supplementary table 6 for a list of the transcript and gene sequences analysed. The online tool Splign (Kapustin et al. 2008) (https://www.ncbi.nlm.nih.gov/sutils/splign/splign.cgi) was employed to determine the exon/intron structure of genes and schematic figures showing gene structure were generated using IBS 1.0 (Liu et al. 2015).

## Acknowledgments

The work reported in this paper was supported by grants from the BBSRC awarded to M.R.E (BB/M001644/1) and A.M.J. (BB/M001032/1). L.A.Y.G was supported by a PhD studentship awarded by the Mexican Council of Science and Technology (CONACyT studentship no. 418612) and Queen Mary University of London and by a Leverhulme Trust grant (RPG-2016-353) awarded to M.R.E. X.Z. was supported by a PhD studentship awarded by the China Scholarship Council and Queen Mary University of London.

## SUPPLEMENTARY FIGURES

**Supplementary Figure 1.**
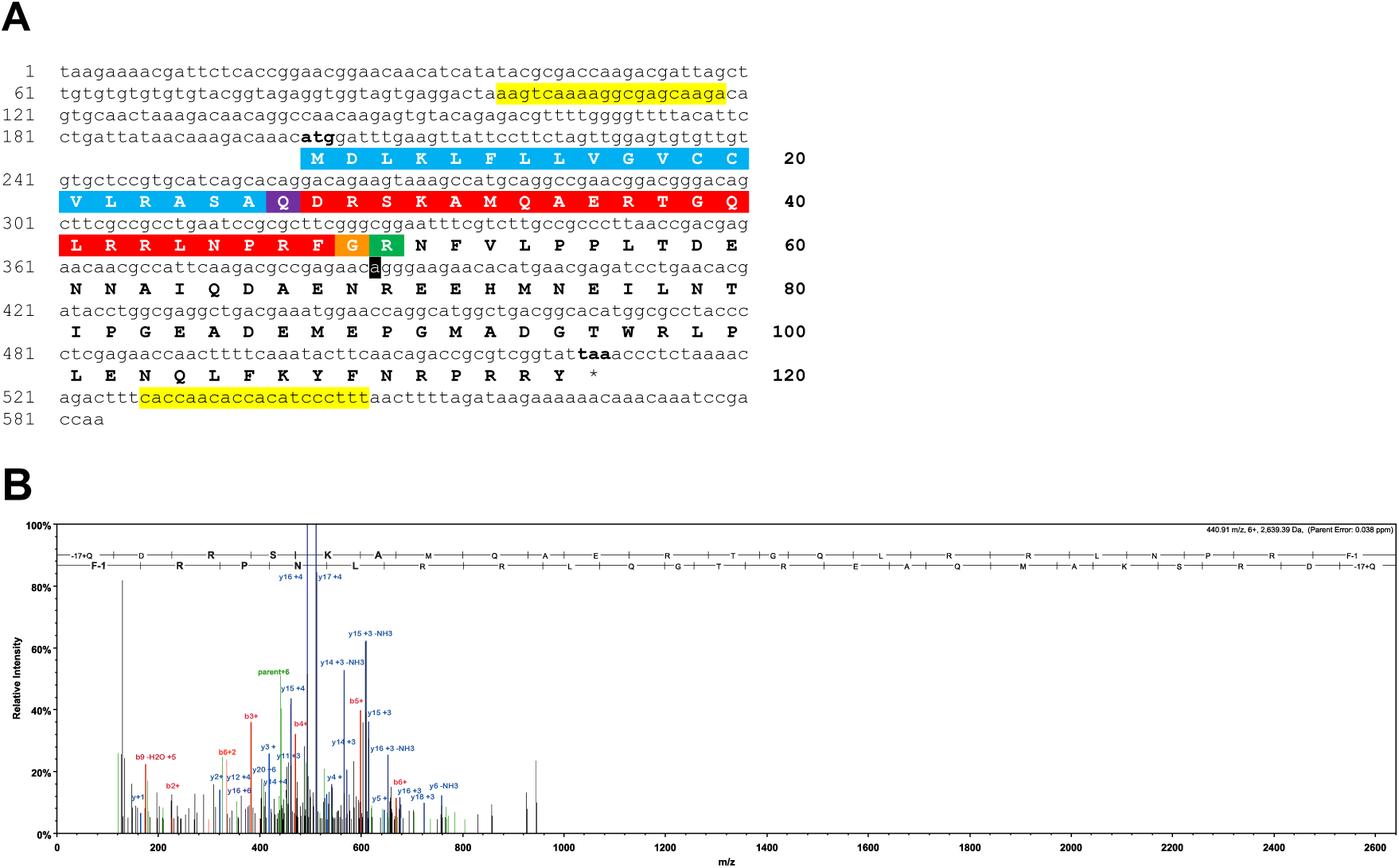
(**A**) Sequence of a cDNA encoding the ArNPLYP precursor. The cDNA sequence (lowercase) comprises an open reading frame of 324 bases that encode a 108-residue protein (uppercase). The predicted signal peptide is shown in blue, the predicted cleavage site is shown in green and the predicted mature peptide is shown in red but with an N-terminal glutamine is a potential substrate for pyroglutamation shown in purple and with a C-terminal glycine is a potential substrate for amidation shown in orange. The cDNA was amplified by PCR from *A. rubens* radial nerve cord cDNA using primers corresponding to the sequences highlighted in yellow. The cDNA was cloned in the vector pBluescript II SK (+) and the T3 and T7 primers were used for the sequencing. A single nucleotide that differs from a contig sequence (1060225) identified from *A. rubens* radial nerve cord transcriptome data is highlighted in black, but this is a synonymous substitution. This sequence has been deposited in GenBank under the accession number MK033631.1 (**B**) Annotated mass spectrum showing the structure of ArNPYL peptide isolated from an *A. rubens* radial nerve cord extract. The peptide QDRSKAMQAERTGQLRRLNPRF, with Q1-Pyroglutamate (−17.02655 Da) and F22-amidated (−0.98402 Da), was observed at charge state +6, monoisotopic m/z 440.90631 Da with a precursor mass error of 0.12 ppm [MH+ 2640.40148 Da] and with a retention time (RT) of 66.3484 min. The b series of peptide fragment ions are shown in red, the y series in blue and additional identified peptide fragment ions in green. The peptide was identified with: Sequest HT (v1.17); XCorr: 4.27, Percolator q-Value: 0.0e0, Percolator PEP:1.6e-2. The fragment match tolerance used for the search was 0.8 Da. The ion table for this mass spectrum can be found in supplementary table 7.

**Supplementary figure 2.**
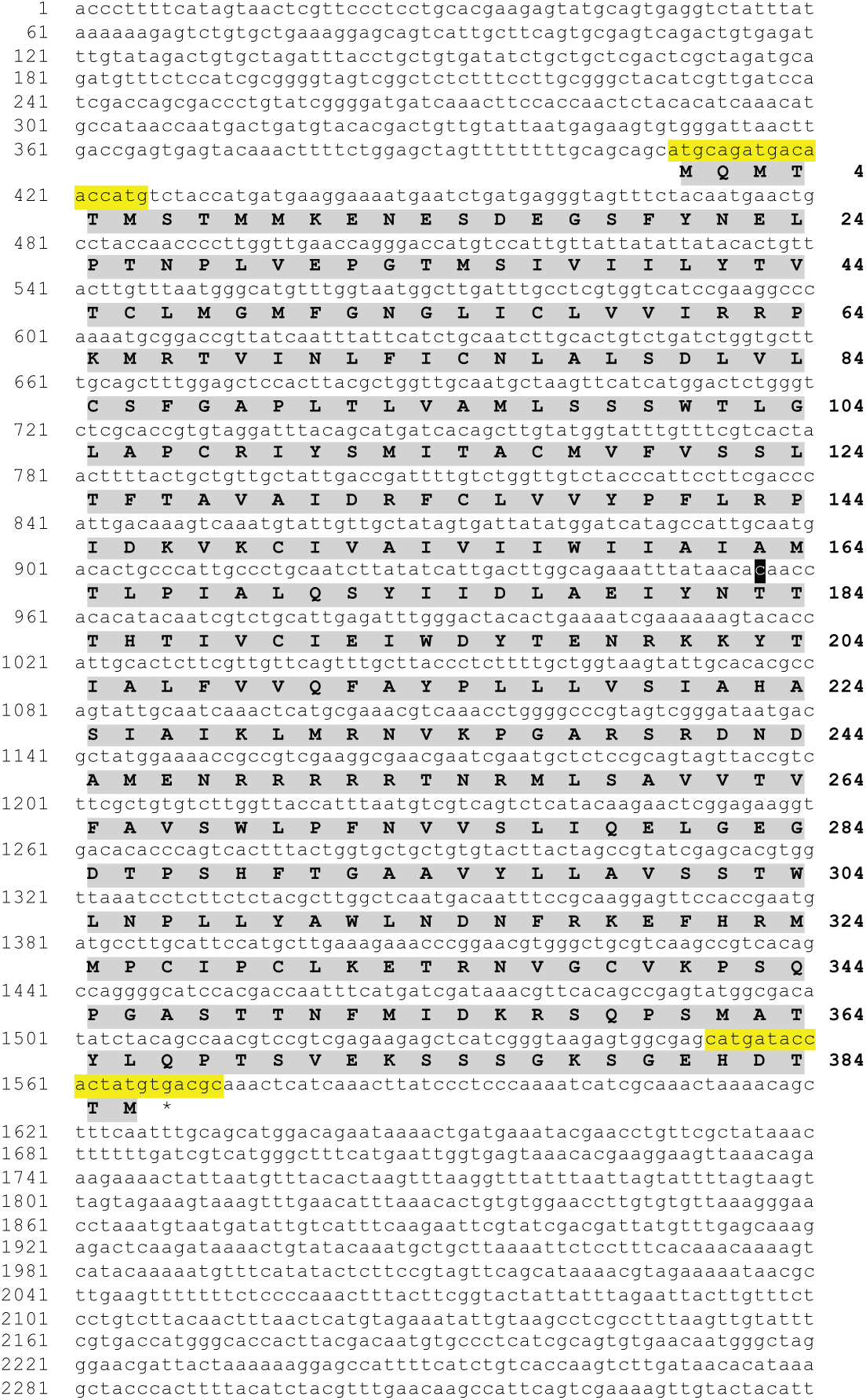
*Asterias rubens* sNPF-type receptor (Ar-sNPFR). Sequence of a transcript (contig 1120879 from *A. rubens* radial nerve cord transcriptome) that encodes a sNPF-type receptor. The transcript sequence (lowercase; numbering on the left) and the deduced amino acid sequence (uppercase and highlighted in grey; numbering on the right) are shown. A cDNA encoding the open reading frame was cloned by PCR using primers corresponding to the sequences highlighted in yellow, ligated into the vector pCDNA 3.1 (+) and sequenced. The sequence of the cloned cDNA was identical to the open reading frame of contig 1120879, with the exception of a single nucleotide substitution (t to c; highlighted in black), which results in replacement of an isoleucine residue (codon: ata in contig 1120879) with a threonine residue (codon: aca in the cloned cDNA). This sequence has been deposited in GenBank under the accession number MH807444.1

**Supplementary Figure 3.**
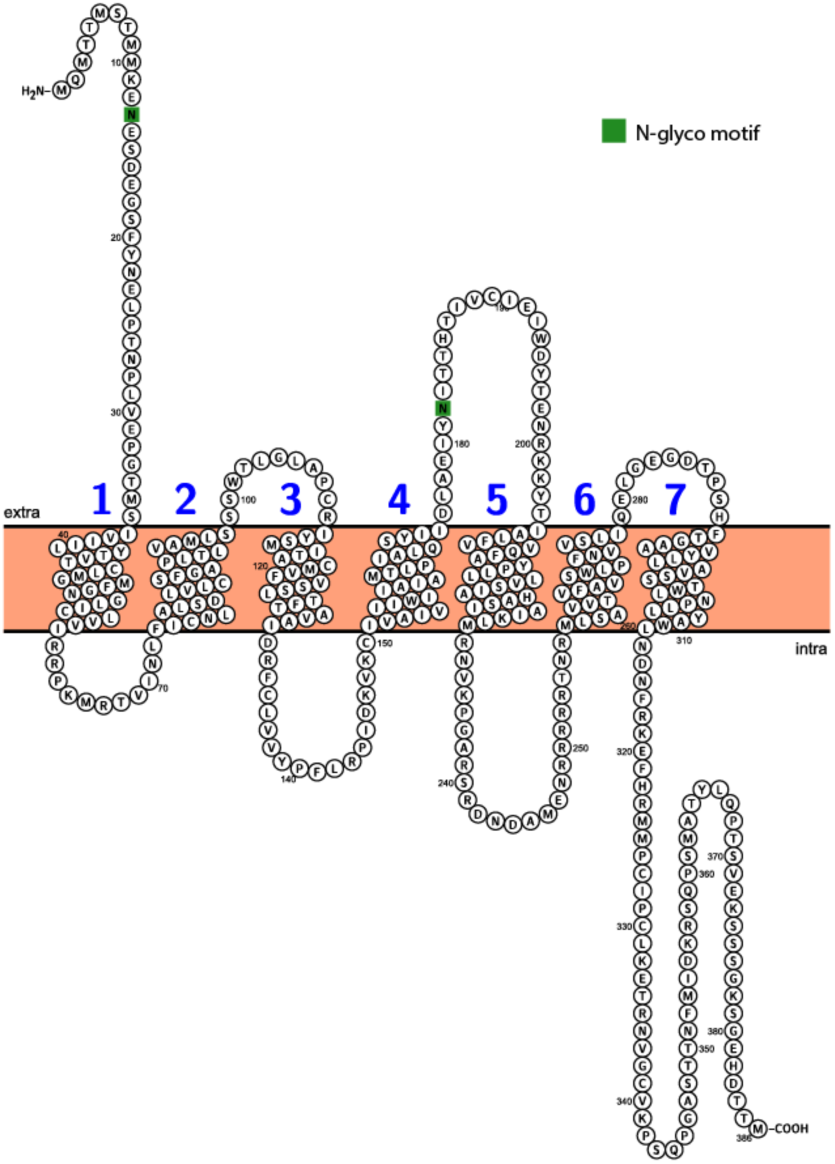
Predicted topology of Ar-sNPFR. Seven predicted transmembrane domains are numbered successively in blue and two predicted N-glycosylation sites are shown with green boxes. This figure was generated using Protter (http://wlab.ethz.ch/protter/start/).

**Supplementary Figure 4.**
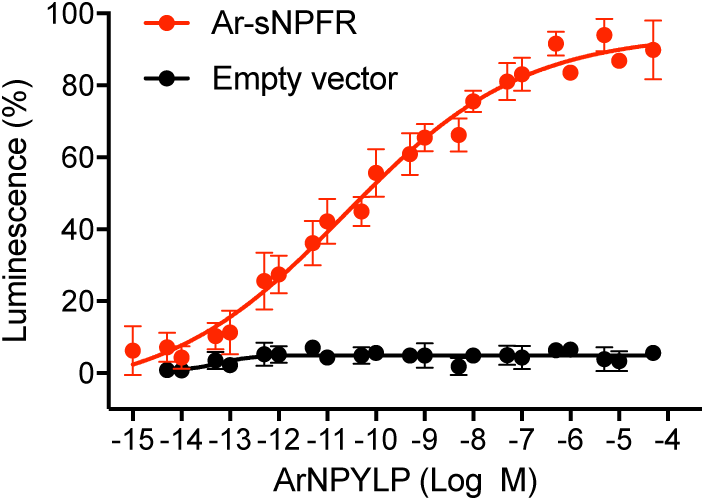
The *A. rubens* NPY-like peptide ArNPYLP does not trigger luminscence in CHO-K1 cells transfected with an empty pcDNA 3.1(+) vector. The peptide ArNPYLP triggers dose-dependent luminscence in CHO-K1 cells transfected with pcDNA 3.1 (+) vector containing the coding sequence for the *A. rubens* sNPY-type receptor Ar-sNPFR (red circles; as also shown in Figure 4) but ArNPYLP does not trigger luminscence in CHO-K1 cells transfected with an empty pcDNA 3.1(+) vector (black circles). Each point represents mean values (± S.E.M.) from at least three independent experiments, with each experiment performed in triplicate.

**Supplementary Figure 5.**
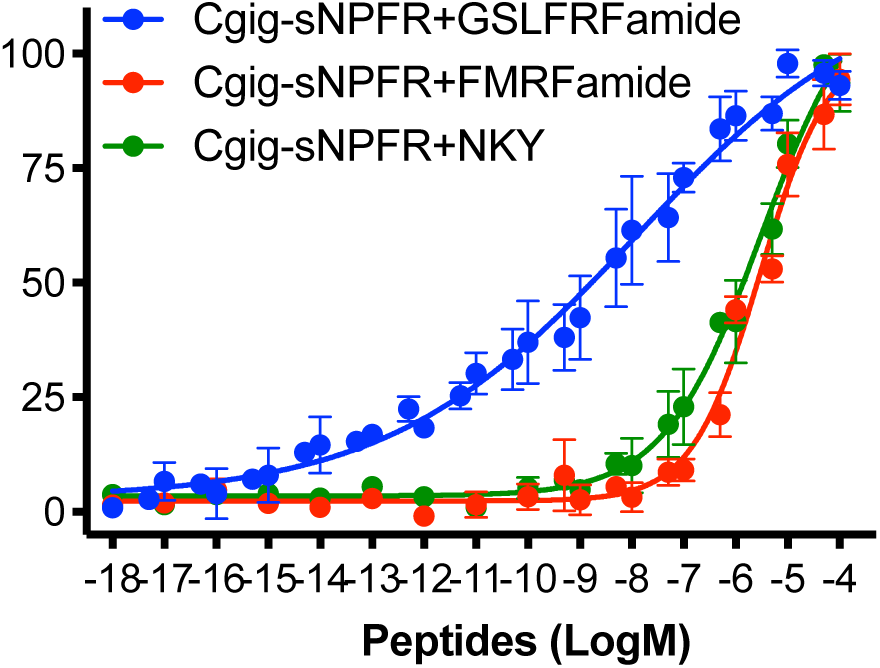
Comparison of three neuropeptides as ligands for the *Crassostrea gigas* sNPF-type receptor (Cgig-sNPFR). The *C. gigas* sNPF-type peptide GSLFRFamide, the *C. gigas* NKY-type peptide Cgig-NKY and the amidated tetrapeptide FMRFamide all cause dose-dependent luminescence in CHO-K1 cells expressing Cgig-sNPFR, the promiscuous G-protein Gα16 and the calcium sensitive luminescent GFP-apoaequorin fusion protein G5A. Each point represents mean values (± S.E.M) from at least three independent experiments done in triplicate. Cgig-NKY and FMRFamide both cause receptor activation but only at relatively high concentrations (EC_50_ values are 3.02 µM and 3.4 µM, respectively), whereas the EC_50_ value for the Cgig-sNPF-type peptide GSLFRFamide is 31 nM. Therefore, GSLFRFamide is likely to act as a ligand for Cgig-sNPFR physiologically, as proposed previously by Bigot et al. (2014). The ability Cgig-NKY and FMRFamide to activate Cgig-sNPFR when heterologously expressed in CHO-K1 cells may reflect non-physiological neuropeptide-receptor crosstalk.

**Methods for supplementary figure 5 –** A cDNA encoding *C. gigas* sNPFR (Bigot et al. 2014) was synthesised, including a 5’ partial Kozak sequence (CACC), by GenScript® (New Jersey, USA) and then cloned into the eukaryotic expression vector pcDNA 3.1(+) (Invitrogen; Cat. No. V790-20). CHO-K1 cells were cultured and transfected with Cgig-sNPFR and then peptide-induced luminescence was measured using methods described previously (Yañez-Guerra et al. 2018). After an incubation period of 3 h with coelenterazine H (Invitrogen Cat. No. C6780), cells were exposed to the synthetic peptides GSLFRFamide, FMRFamide or *C. gigas* NKY (GGIWIWMPAQGYVSVPRDEVGGASNKGSSSNLLRY-NH_2_), all of which were custom synthesised by Peptide Protein Research Ltd. (Fareham, UK). Peptides were diluted in DMEM/F12 Nutrient Mixture medium at concentrations ranging from 10^−14^ M to 10^−4^ M in clear bottom 96-well plates (Sigma-Aldrich; Cat. No. CLS3603-48EA). Responses were normalised and dose-reponse curves were generated as reported in the methodology for the pharmacological characterisation of *A. rubens* sNPF-type receptor Ar-sNPFR (see main methods section of the paper).

## SUPPLEMENTARY TABLES

**Supplementary Table 1.** Accession numbers of the precursor sequences used for the neuropeptide alignment in Figure 1.

| Species name | Accession number or reference |
| --- | --- |
| *Asterias rubens* | QBB78493.1 |
| *Acanthaster planci* | XP_022086679.1 |
| *Amphiura filiformis* | (Zandawala et al. 2017) |
| *Strongylocentrotus purpuratus* | XP_001176371.1 |
| *Saccoglossus kowalevskii* | XP_002741972.1 |
| *Homo sapiens* | NP_000896.1 |
| *Gallus gallus* | NP_990804.1 |
| *Danio rerio* | NP_571149.1 |
| *Branchiostoma floridae* | XP_002609542.1 |
| *Monieza expansa* | MH347240.1 |
| *Schmidtea mediterranea* | ADC84429.1 |
| *Octopus bimaculoides* | XP_014777727.1 |
| *Crassostrea gigas* | XP_011448178.1 |
| *Lymnaea stagnalis* | CAB63265.1 |
| *Helobdella robusta* | XP_009026400.1 |
| *Platynereis dumerilii* | GBZT01002538.1 (predicted from mRNA) |
| *Priapulus caudatus* | XP_014681442.1 |
| *Anopheles gambiae* | XP_315165.3 |
| *Drosophila melanogaster* | XP_001953779 |
| *Stomoxys calcitrans* | XP_013099916.1 |
| *Zootermopsis nevadensis* | XP_021923461.1 |
| *Caenorhabditis elegans* | NP_495111.1 |

**Supplementary Table 2.** Accession numbers of the sequences used for the gene structure analysis in Figure 2.

| Species name | mRNA | Protein | Genome |
| --- | --- | --- | --- |
| *Acanthaster planci* | XM_022230987.1 | XP_022086679.1 | NW_019091356.1 |
| *Strongylocentrotus purpuratus* | XM_001176371.3 | XP_001176371.1 | NW_011971016.1 |
| *Saccoglossus kowalevskii* | XM_002741926.2 | XP_002741972.1 | NW_003156735.1 |
| *Homo sapiens* | NM_000905.4 | NP_000896.1 | Whole genome accessible with NCBI SPLIGN tool |
| *Gallus gallus* | NM_205473.1 | NP_990804.1 | Whole genome accessible with NCBI SPLIGN tool |
| *Danio rerio* | NM_131074.2 | NP_571149.1 | Whole genome accessible with NCBI SPLIGN tool |
| *Branchiostoma floridae* | XM_002609496.1 | XP_002609542.1 | NW_003101541.1 |
| *Octopus bimaculoides* | XM_014922241.1 | XP_014777727.1 | NW_014672493.1 |
| *Crassostrea gigas* | XM_011449876.2 | XP_011448178.1 | NW_011936732.1 |
| *Helobdella robusta* | XM_009028152.1 | XP_009026400.1 | NW_008705278.1 |
| *Priapulus caudatus* | XM_014825956.1 | XP_014681442.1 | NW_014577064.1 |
| *Anopheles gambiae* | XM_315165.4 | XP_315165.3 | NT_078266.2 |
| *Caenorhabditis elegans* | NM_062710.5 | NP_495111.1 | NC_003280.10 |

**Supplementary Table 3.** Accession numbers of the receptor sequences used for the phylogenetic analysis shown in Figure 3.

| Species | Accession number | Receptor type based on findings of this study previous used names in brackets |
| --- | --- | --- |
| *Xenopus tropicalis* | XP_004911210.1 | NPYR2 |
| *Latimeria chalumnae* | XP_014350978.1 | NPYR2 |
| *Homo sapiens* | NP_000901.1 | NPYR2 |
| *Gallus gallus* | NP_001026299.1 | NPYR2 |
| *Saccoglossus kowalevskii* | XP_006817255.1 | NPY/NPF-R |
| *Aplysia californica* | XP_005089627.1, XP_005089880.1 | NPY/NPF-R |
| *Lottia gigantea* | XP_009066442.1 | NPY/NPF-R |
| *Lymnaea stagnalis* | CAA57620.1 | NPY/NPF-R (GPR 105) |
| *Capitela teleta* | ELT88377.1, ELT98787.1 | NPY/NPF-R |
| *Platynereis dumerilii* | AKQ63068.1 | NPY/NPF-R (GPCR 62) |
| *Crassostrea gigas* | XP_011444490.1 | NPY/NPF-R |
| *Schmidtea mediterranea* | ANO39140.1 | NPY/NPF-R (NPY5) |
| *Schmidtea mediterranea* | ANO39130.1 | NPY/NPF-R (NPY1) |
| *Schmidtea mediterranea* | ANO39139.1 | NPY/NPF-R (NPY3) |
| *Schmidtea mediterranea* | ANO39141.1 | NPY/NPF-R (NPY6) |
| *Tribolium castaneum* | XP_008198436.1 | NPY/NPF-R |
| *Drosophila melanogaster* | NP_001246947.1 | NPY/NPF-R |
| *Bactrocera dorsalis* | XP_011202740.1 | NPY/NPF-R |
| *Aedes aegypti* | XP_021693392.1 | NPY/NPF-R |
| *Caenorhabditis elegans* | NP_508234.2 | NPY/NPF-R (NPR-11) |
| *Pristionchus pacificus* | PDM84819.1 | NPY/NPF-R (NPR-11) |
| *Caenorhabditis elegans* | NP_001293732.1 | NPY/NPF-R (NPR-12) |
| *Pristionchus pacificus* | PDM75274.1 | NPY/NPF-R (NPR-12) |
| *Asterias rubens* | MH807444 | sNPFR |
| *Acanthaster planci* | XP_022101544.1 | sNPFR |
| *Strongylocentrotus purpuratus* | XP_003725178.1 | sNPFR |
| *Apostichopus japonicus* | PIK36230.1 | sNPFR |
| *Crassostrea gigas* | XP_011451552.1 | sNPFR |
| *Platynereis dumerilii* | AKQ63001.1 | sNPFR (NKY receptor) |
| *Capitela teleta* | ELT88594.1 | sNPFR |
| *Schmidtea mediterranea* | ANO39142.1 | sNPFR (NPY7) |
| *Schmidtea mediterranea* | ANO39143.1 | sNPFR (NPY8) |
| *Schmidtea mediterranea* | ANO39131.1 | sNPFR (NPY10) |
| *Schmidtea mediterranea* | ANO39144.1 | sNPFR (NPY9) |
| *Drosophila melanogaster* | NP_524176.1 | sNPFR |
| *Tribolium castaneum* | XP_966794.1 | sNPFR |
| *Bombyx mori* | NP_001127707.1 | sNPFR (GPR A10) |
| *Bombyx mori* | NP_001127742.1 | sNPFR (GPR A7) |
| *Bombyx mori* | NP_001127708.1 | sNPFR (GPR A11) |
| *Aedes aegypti* | AGX84998.1 | sNPFR |
| *Priapulus caudatus* | XP_014669822.1 | sNPFR |
| *Caenorhabditis elegans* | NP_508816.1 | sNPFR (NPR1) |
| *Pristionchus pacificus* | PDM70653.1 | sNPFR (NPR1) |
| *Caenorhabditis elegans* | NP_501701.2 | sNPFR (NPR2) |
| *Caenorhabditis elegans* | CAB05681.1 | sNPFR (NPR3) |
| *Caenorhabditis elegans* | NP_001300304.1 | sNPFR (NPR4) |
| *Pristionchus pacificus* | PDM83853.1 | sNPFR (NPR4) |
| *Caenorhabditis elegans* | CCD70460.1 | sNPFR (NPR5) |
| *Pristionchus pacificus* | PDM73363.1 | sNPFR (NPR5) |
| *Homo sapiens* | NP_004239.1 | Prolactin-releasing peptide receptor |
| *Danio rerio* | NP_001034615.1 | Prolactin-releasing peptide receptor |
| *Gallus gallus* | AAW30382.1 | Prolactin-releasing peptide receptor |
| *Alligator mississippiensis* | XP_019341424.1 | Prolactin-releasing peptide receptor |
| *Xenopus tropicalis* | XP_002940396.1 | Prolactin-releasing peptide receptor |
| *Branchistoma belcheri* | XP_019646333.1 | Prolactin-releasing peptide receptor |
| *Branchistoma floridae* | XP_002608333.1 | Prolactin-releasing peptide receptor |
| *Saccoglossus kowalevskii* | XP_002740053.1, | Prolactin-releasing peptide |
|  | XP_006815575.1,<br>XP_002738225.1 | receptor |
| *Homo sapiens* | NP_057624.3 | GPCR83 |
| *Callorhinchus milii* | XP_007900281.1 | GPCR83 |
| *Gallus gallus* | AEO92092.1 | GPCR83 |
| *Takifugus rubripes* | XP_011617071.1 | GPCR83 |
| *Xenopus laevis* | XP_018096593.1 | GPCR83 |
| *Apostichopus japonicus* | PIK61930.1 | GPCR83 |
| *Strongylocentrotus purpuratus* | XP_003729750.1 | GPCR83 |
| *Asterias rubens* | MG744509, MG744510 | Luqin |
| *Strongylocentrotus purpuratus* | XP_783326.1,<br>XP_783390.1 | Luqin |
| *Saccoglossus kowalevskii* | XM_002731957.1,<br>XM_002731958.1,<br>XM_006813011.1,<br>XM_002731956.1 | Luqin |
| *Aplysia californica* | XP_012937781.1 | Luqin |
| *Lottia gigantea* | XP_009064514.1,<br>XP_009064591.1 | Luqin |
| *Lymnea stagnalis* | AAB92258.1 | Luqin |
| *Octopus bimaculoides* | XP_014786450.1 | Luqin |
| *Capitella teleta* | ELT96089.1 | Luqin |
| *Platynereis dumerilii* | KP420214.1 | Luqin |
| *Priapulius caudatus* | XP_014666446.1,<br>XP_014678140.1 | Luqin |
| *Acyrtosiphon pisum* | XP_008178727.1,<br>XP_003241610.1 | RYamide |
| *Tribolium castaneum* | HQ709383.1 | RYamide |
| *Aedes aegypti* | AGX85003.1 | RYamide |
| *Drosophila melanogaster* | P25931.2 | RYamide |
| *Caenorhabditis elegans* | NP_001023541.1 | Luqin |
| *Trichuris suis* | KFD65303.1 | Luqin |
| *Homo sapiens* | AAB20303.1,<br>NP_001049.1,<br>NP_001050.1 | Tachykinin |
| *Ciona intestinalis* | XM_009863501.2 | Tachykinin |
| *Asterias rubens* | MG744511, MG744512 | Tachykinin |
| *Strongylocentrotus purpuratus* | XP_011662258.1 | Tachykinin |
| *Octopus vulgaris* | BAD93354.1 | Tachykinin |
| *Aplysia californica* | XP_012936180.1 | Tachykinin |
| *Lottia gigantea* | XP_009062052.1 | Tachykinin |
| *Capitella teleta* | ELT98449.1 | Tachykinin |
| *Urechis unitinctus* | BAB87199.1 | Tachykinin |
| *Drosophila melanogaster* | FBtr0085507 | Tachykinin |
| *Tribolium castaneum* | XP_008194527.2 | Tachykinin |

**Supplementary Table 4.** Accession numbers of the precursor sequences used for the alignment in figure 5.

| Species name | Accession number |
| --- | --- |
| *Asterias rubens* | QBB78493.1 |
| *Acanthaster planci* | XP_022086679.1 |
| *Amphiura filiformis* | (Zandawala et al. 2017) |
| *Strongylocentrotus purpuratus* | XP_001176371.1 |
| *Crassostrea gigas* | EKC33711.1 |
| *Lymnaea stagnalis* | AAV41057.1 |
| *Platynereis dumerilii* | AEE25645.1 |
| *Schmidtea mediterranea* | DAA33926.1 |
| *Drosophila melanogaster* | NP_724239.1 |
| *Bombyx mori* | NP_001127729.1 |
| *Tribolium castaneum* | DAA34847.1 |
| *Caenorhabditis elegans* flp3 | NP_509694.1 |
| *Caenorhabditis elegans* flp15 | NP_499820.1 |
| *Caenorhabditis elegans* flp18 | NP_508514.2 |
| *Caenorhabditis elegans* flp21 | NP_505011.2 |

**Supplementary Table 5.** Accession numbers of sequences used for the gene structure comparison in figure 6.

| Species name | mRNA | Protein | Genome |
| --- | --- | --- | --- |
| *Acanthaster planci* | XM_022230987.1 | XP_022086679.1 | NW_019091356.1 |
| *Strongylocentrotus purpuratus* | XM_001176371.3 | XP_001176371.1 | Whole genome accessible with NCBI SPLIGN tool |
| *Crassostrea gigas* | FQ665026.1 | EKC33711.1 | JH819141.1 |
| *Pomacea canaliculata* | XM_025228896.1 | XP_025084681.1 | NC_037591.1 |
| *Tribolium castaneum* | XM_008200483.2 | XP_008198705.1 | Whole genome accessible with NCBI SPLIGN tool |
| *Apis mellifera* | XM_003250107.4 | XP_003250155.1 | Whole genome accessible with NCBI SPLIGN tool |
| *Drosophila melanogaster* | NM_165316.2 | NP_724239.1 | Whole genome accessible with NCBI SPLIGN tool |
| *Caenorhabditis elegans* flp-3 | NM_077293.6 | NP_509694.1 | NC_003284.9 |
| *Caenorhabditis elegans* flp-15 | NM_067419.3 | NP_499820.1 | NC_003281.10 |
| *Caenorhabditis elegans* flp-18 | NM_076113.5 | NP_508514.2 | NC_003284.9 |
| *Caenorhabditis elegans* flp-21 | NM_072610.6 | NP_505011.2 | NC_003283.11 |

**Supplementary Table 6.** Accession numbers of sequences used for the alignment and gene structure comparison in figure 7.

| Species name | mRNA | Protein | Genome |
| --- | --- | --- | --- |
| *Asterias rubens* |  | QBB78493.1 |  |
| *Acanthaster planci* | XM_022230987.1 | XP_022086679.1 | NW_019091356.1 |
| *Amphiura filiformis* |  | (Zandawala et al. 2017) |  |
| *Strongylocentrotus purpuratus* | XM_001176371.3 | XP_001176371.1 | Whole genome accessible with NCBI SPLIGN tool |
| *Homo sapiens* | BC069284.1 | AAH69284.1 | Whole genome accessible with NCBI SPLIGN tool |
| *Xenopus laevis* |  | OCT75971.1 |  |
| *Gallus gallus* | NM_001082419.1 | NP_001075888.1 | Whole genome accessible with NCBI SPLIGN tool |
| *Python bivittatus* | XM_025171672.1 | XP_025027440.1 | NW_006532108.1 |
| *Danio rerio* | NM_001245985.1 | NP_001232914.1 | Whole genome accessible with NCBI SPLIGN tool |
| *Latimeria chalumnae* |  | XP_006008888.1 |  |
| *Lepisosteus oculatus* |  | ALD51284.1 |  |
| *Paramormyrops kingsleyae* |  | XP_023653061.1 |  |
| *Petromyzon marinus* |  | ALD51287.1 |  |
| *Branchiostoma floridae* | XM_002595829.1 | XP_002595875.1 | ABEP02025055.1 |
| *Saccoglossus kowalevskii* | XM_002737009.1 | XP_002737055.1 | NW_003134358.1 |
| *Saccoglossus kowalevskii 2* |  | Personal communication | NW_003156735.1 |

**Supplementary Table 7.** Fragmentation table of the mass spectra for the Ar-sNPF peptide (QDRSKAMQAERTGQLRRLNPRF), F22-Amidated (−0.98402 Da), Q1-Pyroglutamate (−17.02655 Da), observed at charge state +6, monoisotopic m/z 440.90631 Da with a precursor mass error of 0.12 ppm).

| #1 | b<sup>+</sup> | b<sup>2+</sup> | b<sup>3+</sup> | b<sup>4+</sup> | b<sup>5+</sup> | b<sup>6+</sup> | Seq. | y<sup>+</sup> | y<sup>2+</sup> | y<sup>3+</sup> | y<sup>4+</sup> | y<sup>5+</sup> | y<sup>6+</sup> | #2 |
| --- | --- | --- | --- | --- | --- | --- | --- | --- | --- | --- | --- | --- | --- | --- |
| 1 | 112.03931 | 56.52329 | 38.01795 | 28.76528 | 23.21368 | 19.51261 | Q-Gln-<br>>pyro-Glu |  |  |  |  |  |  | 22 |
| 2 | 227.06625 | 114.03676 | 76.36027 | 57.52202 | 46.21907 | 38.68377 | D | 2529.36915 | 1265.18821 | 843.79457 | 633.09774 | 506.67965 | 422.40092 | 21 |
| 3 | 383.16736 | 192.08732 | 128.39397 | 96.54730 | 77.43929 | 64.70062 | R | 2414.34221 | 1207.67474 | 805.45225 | 604.34101 | 483.67426 | 403.22976 | 20 |
| 4 | 470.19939 | 235.60333 | 157.40465 | 118.30530 | 94.84570 | 79.20596 | S | 2258.24110 | 1129.62419 | 753.41855 | 565.31573 | 452.45404 | 377.21291 | 19 |
| 5 | 598.29435 | 299.65081 | 200.10297 | 150.32905 | 120.46469 | 100.55512 | K | 2171.20907 | 1086.10817 | 724.40787 | 543.55772 | 435.04763 | 362.70758 | 18 |
| 6 | 669.33146 | 335.16937 | 223.78201 | 168.08832 | 134.67211 | 112.39464 | A | 2043.11410 | 1022.06069 | 681.70955 | 511.53398 | 409.42864 | 341.35841 | 17 |
| 7 | 800.37195 | 400.68961 | 267.46217 | 200.84844 | 160.88021 | 134.23472 | M | 1972.07699 | 986.54213 | 658.03051 | 493.77471 | 395.22122 | 329.51890 | 16 |
| 8 | 928.43053 | 464.71890 | 310.14836 | 232.86309 | 186.49193 | 155.57782 | Q | 1841.03651 | 921.02189 | 614.35035 | 461.01458 | 369.01312 | 307.67881 | 15 |
| 9 | 999.46764 | 500.23746 | 333.82740 | 250.62237 | 200.69935 | 167.41734 | A | 1712.97793 | 856.99260 | 571.66416 | 428.99994 | 343.40141 | 286.33572 | 14 |
| 10 | 1128.51023 | 564.75875 | 376.84160 | 282.88302 | 226.50787 | 188.92444 | E | 1641.94081 | 821.47405 | 547.98512 | 411.24066 | 329.19398 | 274.49620 | 13 |
| 11 | 1284.61134 | 642.80931 | 428.87530 | 321.90829 | 257.72809 | 214.94129 | R | 1512.89822 | 756.95275 | 504.97092 | 378.98001 | 303.38547 | 252.98910 | 12 |
| 12 | 1385.65902 | 693.33315 | 462.55786 | 347.17021 | 277.93763 | 231.78257 | T | 1356.79711 | 678.90219 | 452.93722 | 339.95474 | 272.16524 | 226.97225 | 11 |
| 13 | 1442.68049 | 721.84388 | 481.56501 | 361.42558 | 289.34192 | 241.28614 | G | 1255.74943 | 628.37835 | 419.25466 | 314.69282 | 251.95571 | 210.13097 | 10 |
| 14 | 1570.73906 | 785.87317 | 524.25121 | 393.44022 | 314.95363 | 262.62924 | Q | 1198.72797 | 599.86762 | 400.24751 | 300.43745 | 240.55142 | 200.62739 | 9 |
| 15 | 1683.82313 | 842.41520 | 561.94589 | 421.71124 | 337.57045 | 281.47659 | L | 1070.66939 | 535.83833 | 357.56131 | 268.42281 | 214.93970 | 179.28430 | 8 |
| 16 | 1839.92424 | 920.46576 | 613.97960 | 460.73652 | 368.79067 | 307.49344 | R | 957.58533 | 479.29630 | 319.86663 | 240.15179 | 192.32289 | 160.43695 | 7 |
| 17 | 1996.02535 | 998.51631 | 666.01330 | 499.76179 | 400.01089 | 333.51029 | R | 801.48422 | 401.24575 | 267.83292 | 201.12651 | 161.10266 | 134.42010 | 6 |
| 18 | 2109.10941 | 1055.05834 | 703.70799 | 528.03281 | 422.62770 | 352.35763 | L | 645.38311 | 323.19519 | 215.79922 | 162.10123 | 129.88244 | 108.40325 | 5 |
| 19 | 2223.15234 | 1112.07981 | 741.72230 | 556.54354 | 445.43629 | 371.36479 | N | 532.29904 | 266.65316 | 178.10453 | 133.83022 | 107.26563 | 89.55590 | 4 |
| 20 | 2320.20510 | 1160.60619 | 774.07322 | 580.80673 | 464.84684 | 387.54025 | P | 418.25611 | 209.63170 | 140.09022 | 105.31949 | 84.45704 | 70.54875 | 3 |
| 21 | 2476.30622 | 1238.65675 | 826.10692 | 619.83201 | 496.06706 | 413.55710 | R | 321.20335 | 161.10531 | 107.73930 | 81.05630 | 65.04649 | 54.37329 | 2 |
| 22 |  |  |  |  |  |  | F-amidated | 165.10224 | 83.05476 | 55.70560 | 42.03102 | 33.82627 | 28.35644 | 1 |

## Bibliography

Adrian TE, Allen JM, Bloom SR, Ghatei MA, Rossor MN, Roberts GW, Crow TJ, Tatemoto K, Polak JM. 1983. Neuropeptide Y distribution in human brain. Nature 306:584– 586.

Ament SA, Velarde RA, Kolodkin MH, Moyse D, Robinson GE. 2011. Neuropeptide Y-like signalling and nutritionally mediated gene expression and behaviour in the honey bee. Insect Mol Biol 20:335–345.

Bard JA, Walker MW, Branchek TA, Weinshank RL. 1995. Cloning and functional expression of a human Y4 subtype receptor for pancreatic polypeptide, neuropeptide Y, and peptide YY. J Biol Chem 270:26762–26765.

Baubet V, Le Mouellic H, Campbell AK, Lucas-Meunier E, Fossier P, Brúlet P. 2000. Chimeric green fluorescent protein-aequorin as bioluminescent Ca2+ reporters at the single-cell level. Proc Natl Acad Sci U S A 97:7260–7265.

Bauknecht P, Jékely G. 2015. Large-Scale Combinatorial Deorphanization of Platynereis Neuropeptide GPCRs. Cell Rep 12:684–693.

Bigot L, Beets I, Dubos M-P, Boudry P, Schoofs L, Favrel P. 2014. Functional characterization of a short neuropeptide F-related receptor in a lophotrochozoan, the mollusk Crassostrea gigas. J Exp Biol 217:2974–2982.

Blomqvist AG, Söderberg C, Lundell I, Milner RJ, Larhammar D. 1992. Strong evolutionary conservation of neuropeptide Y: sequences of chicken, goldfish, and Torpedo marmorata DNA clones. Proc Natl Acad Sci U S A 89:2350–2354.

Blundell TL, Pitts JE, Tickle IJ, Wood SP, Wu CW. 1981. X-ray analysis (1. 4-A resolution) of avian pancreatic polypeptide: Small globular protein hormone. Proc Natl Acad Sci U S A 78:4175–4179.

Brockmann A, Annangudi SP, Richmond TA, Ament SA, Xie F, Southey BR, Rodriguez-Zas SR, Robinson GE, Sweedler JV. 2009. Quantitative peptidomics reveal brain peptide signatures of behavior. Proc Natl Acad Sci U S A 106:2383–2388.

Brown MR, Crim JW, Arata RC, Cai HN, Chun C, Shen P. 1999. Identification of a Drosophila brain-gut peptide related to the neuropeptide Y family. Peptides 20:1035– 1042.

Burge CB, Karlin S. 1998. Finding the genes in genomic DNA. Curr Opin Struct Biol 8:346– 354.

Burge C, Karlin S. 1997. Prediction of complete gene structures in human genomic DNA. J Mol Biol 268:78–94.

Cardoso JCR, Félix RC, Fonseca VG, Power DM. 2012. Feeding and the rhodopsin family g-protein coupled receptors in nematodes and arthropods. Front Endocrinol (Lausanne) 3:157.

Cerdá-Reverter JM, Martínez-Rodríguez G, Zanuy S, Carrillo M, Larhammar D. 2000. Molecular evolution of the neuropeptide Y (NPY) family of peptides: cloning of three NPY-related peptides from the sea bass (Dicentrarchus labrax). Regul Pept 95:25–34.

Chalasani SH, Kato S, Albrecht DR, Nakagawa T, Abbott LF, Bargmann CI. 2010. Neuropeptide feedback modifies odor-evoked dynamics in Caenorhabditis elegans olfactory neurons. Nat Neurosci 13:615–621.

Chen M-E, Pietrantonio PV. 2006. The short neuropeptide F-like receptor from the red imported fire ant, Solenopsis invicta Buren (Hymenoptera: Formicidae). Arch Insect Biochem Physiol 61:195–208.

Christ P, Hill SR, Schachtner J, Hauser F, Ignell R. 2018. Functional characterization of mosquito short neuropeptide F receptors. Peptides 103:31–39.

Cohen M, Reale V, Olofsson B, Knights A, Evans P, de Bono M. 2009. Coordinated regulation of foraging and metabolism in C. elegans by RFamide neuropeptide signaling. Cell Metab 9:375–385.

Conzelmann M, Williams EA, Krug K, Franz-Wachtel M, Macek B, Jékely G. 2013. The neuropeptide complement of the marine annelid Platynereis dumerilii. BMC Genomics 14:906.

Curry WJ, Shaw C, Johnston CF, Thim L, Buchanan KD. 1992. Neuropeptide F: primary structure from the tubellarian, Artioposthia triangulata. Comp Biochem Physiol C, Comp Pharmacol Toxicol 101:269–274.

Díaz-Miranda L, Escalona de Motta G, García-Arrarás JE. 1991. Localization of neuropeptides in the nervous system of the marine annelid Sabellastarte magnifica. Cell Tissue Res 266:209–217.

Dillen S, Verdonck R, Zels S, Van Wielendaele P, Broeck JV. 2013. Identification of the short Neuropeptide F precursor in the desert locust: Evidence for an inhibitory role of sNPF in the control of feeding. Peptides.

Dillen S, Zels S, Verlinden H, Spit J, Van Wielendaele P, Vanden Broeck J. 2013. Functional characterization of the short neuropeptide F receptor in the desert locust, Schistocerca gregaria. PLoS ONE 8:e53604.

Edgar RC. 2004. MUSCLE: multiple sequence alignment with high accuracy and high throughput. Nucleic Acids Res 32:1792–1797.

Elphick MR, Mirabeau O. 2014. The Evolution and Variety of RFamide-Type Neuropeptides: Insights from Deuterostomian Invertebrates. Front Endocrinol (Lausanne) 5:93.

Elphick MR, Mirabeau O, Larhammar D. 2018. Evolution of neuropeptide signalling systems. J Exp Biol 221.

Ezcurra M, Walker DS, Beets I, Swoboda P, Schafer WR. 2016. Neuropeptidergic Signaling and Active Feeding State Inhibit Nociception in Caenorhabditis elegans. J Neurosci 36:3157–3169.

Fadda M, Hasakiogullari I, Temmerman L, Beets I, Zels S, Schoofs L. 2019. Regulation of feeding and metabolism by neuropeptide F and short neuropeptide F in invertebrates. Front Endocrinol (Lausanne) 10:64.

Farzi A, Reichmann F, Holzer P. 2015. The homeostatic role of neuropeptide Y in immune function and its impact on mood and behaviour. Acta Physiol (Oxf) 213:603–627.

Gabaldón T. 2008. Large-scale assignment of orthology: back to phylogenetics? Genome Biol 9:235.

Garczynski SF, Brown MR, Shen P, Murray TF, Crim JW. 2002. Characterization of a functional neuropeptide F receptor from Drosophila melanogaster. Peptides 23:773– 780.

Garczynski SF, Crim JW, Brown MR. 2007. Characterization and expression of the short neuropeptide F receptor in the African malaria mosquito, Anopheles gambiae. Peptides 28:109–118.

Glover ID, Barlow DJ, Pitts JE, Wood SP, Tickle IJ, Blundell TL, Tatemoto K, Kimmel JR, Wollmer A, Strassburger W. 1984. Conformational studies on the pancreatic polypeptide hormone family. Eur J Biochem 142:379–385.

Glover I, Haneef I, Pitts J, Wood S, Moss D, Tickle I, Blundell T. 1983. Conformational flexibility in a small globular hormone: x-ray analysis of avian pancreatic polypeptide at 0.98-A resolution. Biopolymers 22:293–304.

Hinuma S, Habata Y, Fujii R, Kawamata Y, Hosoya M, Fukusumi S, Kitada C, Masuo Y, Asano T, Matsumoto H, et al. 1998. A prolactin-releasing peptide in the brain. Nature 393:272–276.

Hoek RM, Li KW, van Minnen J, Lodder JC, de Jong-Brink M, Smit AB, van Kesteren RE. 2005. LFRFamides: a novel family of parasitation-induced-RFamide neuropeptides that inhibit the activity of neuroendocrine cells in Lymnaea stagnalis. J Neurochem 92:1073–1080.

Holzer P, Reichmann F, Farzi A. 2012. Neuropeptide Y, peptide YY and pancreatic polypeptide in the gut-brain axis. Neuropeptides 46:261–274.

Jékely G. 2013. Global view of the evolution and diversity of metazoan neuropeptide signaling. Proc Natl Acad Sci U S A 110:8702–8707.

Jékely G, Melzer S, Beets I, Kadow ICG, Koene J, Haddad S, Holden-Dye L. 2018. The long and the short of it - a perspective on peptidergic regulation of circuits and behaviour. J Exp Biol 221.

Jiang H-B, Gui S-H, Xu L, Pei Y-X, Smagghe G, Wang J-J. 2017. The short neuropeptide F modulates olfactory sensitivity of Bactrocera dorsalis upon starvation. J Insect Physiol 99:78–85.

Kapustin Y, Souvorov A, Tatusova T, Lipman D. 2008. Splign: algorithms for computing spliced alignments with identification of paralogs. Biol Direct 3:20.

Kawada T, Sekiguchi T, Sakai T, Aoyama M, Satake H. 2010. Neuropeptides, hormone peptides, and their receptors in Ciona intestinalis: an update. Zool Sci 27:134–153.

Keire DA, Mannon P, Kobayashi M, Walsh JH, Solomon TE, Reeve JR. 2000. Primary structures of PYY, [Pro(34)]PYY, and PYY-(3-36) confer different conformations and receptor selectivity. Am J Physiol Gastrointest Liver Physiol 279:G126–G131.

Kim KM, Sung S, Caetano-Anollés G, Han JY, Kim H. 2008. An approach of orthology detection from homologous sequences under minimum evolution. Nucleic Acids Res 36:e110.

Kubiak TM, Larsen MJ, Bowman JW, Geary TG, Lowery DE. 2008. FMRFamide-like peptides encoded on the flp-18 precursor gene activate two isoforms of the orphan Caenorhabditis elegans G-protein-coupled receptor Y58G8A.4 heterologously expressed in mammalian cells. Biopolymers 90:339–348.

Kubiak TM, Larsen MJ, Nulf SC, Zantello MR, Burton KJ, Bowman JW, Modric T, Lowery DE. 2003. Differential activation of “social” and “solitary” variants of the Caenorhabditis elegans G protein-coupled receptor NPR-1 by its cognate ligand AF9. J Biol Chem 278:33724–33729.

Kubiak TM, Larsen MJ, Zantello MR, Bowman JW, Nulf SC, Lowery DE. 2003. Functional annotation of the putative orphan Caenorhabditis elegans G-protein-coupled receptor C10C6.2 as a FLP15 peptide receptor. J Biol Chem 278:42115–42120.

Kumar S, Stecher G, Tamura K. 2016. MEGA7: molecular evolutionary genetics analysis version 7.0 for bigger datasets. Mol Biol Evol 33:1870–1874.

Lagerström MC, Fredriksson R, Bjarnadóttir TK, Fridmanis D, Holmquist T, Andersson J, Yan Y-L, Raudsepp T, Zoorob R, Kukkonen JP, et al. 2005. Origin of the prolactin-releasing hormone (PRLH) receptors: evidence of coevolution between PRLH and a redundant neuropeptide Y receptor during vertebrate evolution. Genomics 85:688– 703.

Larhammar D. 1996. Evolution of neuropeptide Y, peptide YY and pancreatic polypeptide. Regul Pept 62:1–11.

Larhammar D, Blomqvist AG, Söderberg C. 1993. Evolution of neuropeptide Y and its related peptides. Comp Biochem Physiol C, Comp Pharmacol Toxicol 106:743–752.

Lee K-S, You K-H, Choo J-K, Han Y-M, Yu K. 2004. Drosophila short neuropeptide F regulates food intake and body size. J Biol Chem 279:50781–50789.

Leung PS, Shaw C, Maule AG, Thim L, Johnston CF, Irvine GB. 1992. The primary structure of neuropeptide F (NPF) from the garden snail, Helix aspersa. Regul Pept 41:71–81.

Lin M, Mita M, Egertová M, Zampronio CG, Jones AM, Elphick MR. 2017. Cellular localization of relaxin-like gonad-stimulating peptide expression in Asterias rubens: New insights into neurohormonal control of spawning in starfish. J Comp Neurol 525:1599–1617.

Liu W, Xie Y, Ma J, Luo X, Nie P, Zuo Z, Lahrmann U, Zhao Q, Zheng Y, Zhao Y, et al. 2015. IBS: an illustrator for the presentation and visualization of biological sequences. Bioinformatics 31:3359–3361.

Lundell I, Berglund MM, Starbäck P, Salaneck E, Gehlert DR, Larhammar D. 1997. Cloning and characterization of a novel neuropeptide Y receptor subtype in the zebrafish. DNA Cell Biol 16:1357–1363.

Mair GR, Halton DW, Shaw C, Maule AG. 2000. The neuropeptide F (NPF) encoding gene from the cestode, Moniezia expansa. Parasitology 120:71–77.

Ma Q, Cao Z, Yu Y, Yan L, Zhang W, Shi Y, Zhou N, Huang H. 2017. Bombyx neuropeptide G protein-coupled receptor A7 is the third cognate receptor for short neuropeptide F from silkworm. J Biol Chem 292:20599–20612.

Maule AG, Shaw C, Halton DW, Thim L, Johnston CF, Fairweather I, Buchanan KD. 1991. Neuropeptide F: a novel parasitic flatworm regulatory peptide from Moniezia expansa (Cestoda: Cyclophyllidea). Parasitology 102:309–316.

Mertens I, Meeusen T, Huybrechts R, De Loof A, Schoofs L. 2002. Characterization of the short neuropeptide F receptor from Drosophila melanogaster. Biochem Biophys Res Commun 297:1140–1148.

Minor RK, Chang JW, de Cabo R. 2009. Hungry for life: How the arcuate nucleus and neuropeptide Y may play a critical role in mediating the benefits of calorie restriction. Mol Cell Endocrinol 299:79–88.

Mirabeau O, Joly J-S. 2013. Molecular evolution of peptidergic signaling systems in bilaterians. Proc Natl Acad Sci U S A 110:E2028–E2037.

Morris BJ. 1989. Neuronal localisation of neuropeptide Y gene expression in rat brain. J Comp Neurol 290:358–368.

Nässel DR, Wegener C. 2011. A comparative review of short and long neuropeptide F signaling in invertebrates: Any similarities to vertebrate neuropeptide Y signaling? Peptides 32:1335–1355.

Nygaard R, Nielbo S, Schwartz TW, Poulsen FM. 2006. The PP-fold solution structure of human polypeptide YY and human PYY3-36 as determined by NMR. Biochemistry 45:8350–8357.

Omasits U, Ahrens CH, Müller S, Wollscheid B. 2014. Protter: interactive protein feature visualization and integration with experimental proteomic data. Bioinformatics 30:884–886.

Pedrazzini T, Pralong F, Grouzmann E. 2003. Neuropeptide Y: the universal soldier. Cell Mol Life Sci 60:350–377.

Priyam A, Woodcroft BJ, Rai V, Munagala A, Moghul I, Ter F, Gibbins MA, Moon H, Leonard G, Rumpf W, et al. 2015. Sequenceserver: a modern graphical user interface for custom BLAST databases. BioRxiv.

Putnam NH, Butts T, Ferrier DEK, Furlong RF, Hellsten U, Kawashima T, Robinson-Rechavi M, Shoguchi E, Terry A, Yu J-K, et al. 2008. The amphioxus genome and the evolution of the chordate karyotype. Nature 453:1064–1071.

Rajpara SM, Garcia PD, Roberts R, Eliassen JC, Owens DF, Maltby D, Myers RM, Mayeri E. 1992. Identification and molecular cloning of a neuropeptide Y homolog that produces prolonged inhibition in Aplysia neurons. Neuron 9:505–513.

Roch GJ, Tello JA, Sherwood NM. 2014. At the transition from invertebrates to vertebrates, a novel GnRH-like peptide emerges in amphioxus. Mol Biol Evol 31:765–778.

Rogers C, Reale V, Kim K, Chatwin H, Li C, Evans P, de Bono M. 2003. Inhibition of Caenorhabditis elegans social feeding by FMRFamide-related peptide activation of NPR-1. Nat Neurosci 6:1178–1185.

Saberi A, Jamal A, Beets I, Schoofs L, Newmark PA. 2016. Gpcrs direct germline development and somatic gonad function in planarians. PLoS Biol 14:e1002457.

Satoh N, Rokhsar D, Nishikawa T. 2014. Chordate evolution and the three-phylum system. Proc Biol Sci 281:20141729.

Schoofs L, Clynen E, Cerstiaens A, Baggerman G, Wei Z, Vercammen T, Nachman R, De Loof A, Tanaka S. 2001. Newly discovered functions for some myotropic neuropeptides in locusts. Peptides 22:219–227.

Schwartz TW, Fuhlendorff J, Kjems LL, Kristensen MS, Vervelde M, O’Hare M, Krstenansky JL, Bjørnholm B. 1990. Signal Epitopes in the Three-Dimensional Structure of Neuropeptide Y. Ann N Y Acad Sci 611:35–47.

Semmens DC, Beets I, Rowe ML, Blowes LM, Oliveri P, Elphick MR. 2015. Discovery of sea urchin NGFFFamide receptor unites a bilaterian neuropeptide family. Open Biol 5:150030.

Semmens DC, Mirabeau O, Moghul I, Pancholi MR, Wurm Y, Elphick MR. 2016. Transcriptomic identification of starfish neuropeptide precursors yields new insights into neuropeptide evolution. Open Biol 6:150224.

Simakov O, Kawashima T, Marlétaz F, Jenkins J, Koyanagi R, Mitros T, Hisata K, Bredeson J, Shoguchi E, Gyoja F, et al. 2015. Hemichordate genomes and deuterostome origins. Nature 527:459–465.

Spittaels K, Verhaert P, Shaw C, Johnston RN, Devreese B, Van Beeumen J, De Loof A. 1996. Insect neuropeptide F (NPF)-related peptides: isolation from Colorado potato beetle (Leptinotarsa decemlineata) brain. Insect Biochem Mol Biol 26:375–382.

Tachibana T, Sakamoto T. 2014. Functions of two distinct “prolactin-releasing peptides” evolved from a common ancestral gene. Front Endocrinol (Lausanne) 5:170.

Tatemoto K. 1982. Neuropeptide Y: complete amino acid sequence of the brain peptide. Proc Natl Acad Sci U S A 79:5485–5489.

Tatemoto K, Carlquist M, Mutt V. 1982. Neuropeptide Y--a novel brain peptide with structural similarities to peptide YY and pancreatic polypeptide. Nature 296:659–660.

Tian S, Zandawala M, Beets I, Baytemur E, Slade SE, Scrivens JH, Elphick MR. 2016. Urbilaterian origin of paralogous GnRH and corazonin neuropeptide signalling pathways. Sci Rep 6:28788.

Trifinopoulos J, Nguyen L-T, Haeseler A von, Minh BQ. 2016. W-IQ-TREE: a fast online phylogenetic tool for maximum likelihood analysis. Nucleic Acids Res 44:W232– W235.

Vanden Broeck J. 2001. Neuropeptides and their precursors in the fruitfly, Drosophila melanogaster. Peptides 22:241–254.

Veenstra JA. 2011. Neuropeptide evolution: neurohormones and neuropeptides predicted from the genomes of Capitella teleta and Helobdella robusta. Gen Comp Endocrinol 171:160–175.

Yamanaka N, Yamamoto S, Zitnan D, Watanabe K, Kawada T, Satake H, Kaneko Y, Hiruma K, Tanaka Y, Shinoda T, et al. 2008. Neuropeptide receptor transcriptome reveals unidentified neuroendocrine pathways. PLoS ONE 3:e3048.

Yañez-Guerra LA, Delroisse J, Barreiro-Iglesias A, Slade SE, Scrivens JH, Elphick MR. 2018. Discovery and functional characterisation of a luqin-type neuropeptide signalling system in a deuterostome. Sci Rep 8:7220.

Zandawala M, Moghul I, Yañez Guerra LA, Delroisse J, Abylkassimova N, Hugall AF, O’Hara TD, Elphick MR. 2017. Discovery of novel representatives of bilaterian neuropeptide families and reconstruction of neuropeptide precursor evolution in ophiuroid echinoderms. Open Biol 7.

Zatylny-Gaudin C, Bernay B, Zanuttini B, Leprince J, Vaudry H, Henry J. 2010. Characterization of a novel LFRFamide neuropeptide in the cephalopod Sepia officinalis. Peptides 31:207–214.

Zhang G, Fang X, Guo X, Li L, Luo R, Xu F, Yang P, Zhang L, Wang X, Qi H, et al. 2012. The oyster genome reveals stress adaptation and complexity of shell formation. Nature 490:49–54.

Zhang L, Bijker MS, Herzog H. 2011. The neuropeptide Y system: pathophysiological and therapeutic implications in obesity and cancer. Pharmacol Ther 131:91–113.

